# Uncertainty quantification, propagation and characterization by Bayesian analysis combined with global sensitivity analysis applied to dynamical intracellular pathway models

**DOI:** 10.1101/294884

**Authors:** Olivia Eriksson, Alexandra Jauhiainen, Sara Maad Sasane, Andrei Kramer, Anu G Nair, Carolina Sartorius, Jeanette Hellgren Kotaleski

## Abstract

**Motivation:** Dynamical models describing intracellular phenomena are increasing in size and complexity as more information is obtained from experiments. These models are often over-parameterized with respect to the quantitative data used for parameter estimation, resulting in uncertainty in the individual parameter estimates as well as in the predictions made from the model. Here we combine Bayesian analysis with global sensitivity analysis in order to give better informed predictions; to point out weaker parts of the model that are important targets for further experiments, as well as give guidance on parameters that are essential in distinguishing different qualitative output behaviours.

**Results:** We used approximate Bayesian computation (ABC) to estimate the model parameters from experimental data, as well as to quantify the uncertainty in this estimation (inverse uncertainty quantification), resulting in a *posterior distribution* for the parameters. This parameter uncertainty was next propagated to a corresponding uncertainty in the predictions (forward uncertainty propagation), and a global sensitivity analysis was performed on the prediction using the posterior distribution as the possible values for the parameters. This methodology was applied on a relatively large and complex model relevant for synaptic plasticity, using experimental data from several sources. We could hereby point out those parameters that by themselves have the largest contribution to the uncertainty of the prediction as well as identify parameters important to separate between qualitatively different predictions.This approach is useful both for experimental design as well as model building.

## 1 Introduction

Dynamical models describing intracellular phenomena, like the protein interactions of signalling pathways, are increasing in size and complexity as more information from experiments is incorporated. These models are built from qualitative knowledge about the interaction topology, inferred from experiments like e.g. gene knock-outs, as well as from experimental quantitative data describing the input-output relationship of the observed system (Le Novere, 2015). The quantitative data are often sparse as compared to the size of the system, and trying to estimate parameters based on this data often results in large uncertainty in the parameter values, or that some parameters cannot be constrained at all given the data and model (i.e. are unidentifiable (Raue *et al*., 2009)). Parameter estimation from data (model calibration) rarely leads to precise point estimates for the parameters. Rather, the calibration often gives possible ranges for the parameters, and hence it is useful to provide distributions for the parameters, rather than to focus on single point estimates, i.e. to quantify the uncertainty in the parameter estimates (Vanlier *et al*., 2013). Of interest is also to investigate how the uncertainty in the model parameters are transferred into uncertainty for predictions from the model, and to study how this uncertainty in the predictions can be mapped back and attributed to the different model parameters.

**Fig. 1.**
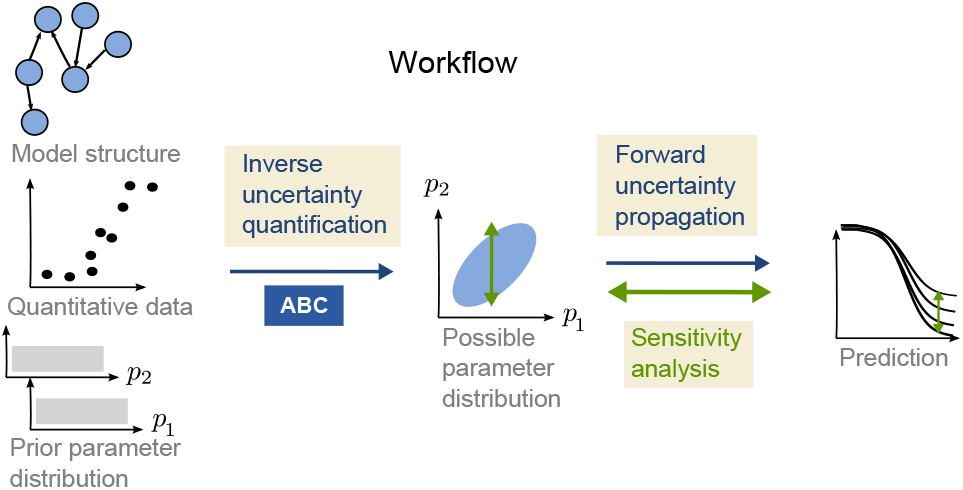
An illustration showing the different parts of the workflow and how global sensitivity analysis is applied on the prediction using the posterior distribution as a restriction on possible model parameter values.

In this paper we develop and combine established methods for Bayesian inference and global sensitivity analysis to show that they, when applied together to a relatively large and complex dynamical system involved in synaptic plasticity, give a comprehensive evaluation of the system given an experimental context, and can guide further experiments and modelling. Uncertainty analysis and global sensitivity analysis have often been performed as separate methods in different modelling studies, but here they are combined so that the global sensitivity analysis is performed based on the posterior distribution of the parameters and we consider system behaviors for which we have no data (i.e. predictions). The sensitivity analysis thereby reveals which parts of the model that are most unconstrained given a certain prediction. We can also compare different hypotheses and draw conclusions about parameters important for a certain model output.

### 1.1 Problem statement

We start from a mathematical model, experimental data and a prior distribution of the parameters describing the prior knowledge (if any), see Figure 1. The model is described by the nonlinear system:

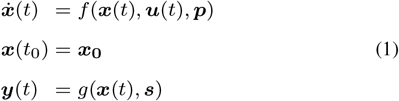

where ***x***(*t*) corresponds to internal state variables (like protein concentrations in an intracellular model), ***u***(*t*) to external input (e.g. an external signal to the cell, or the total amount of a specific protein), ***y***(*t*) are the outputs, i.e. the observed variables (modelling counterparts to possible experimental readouts), ***p*** are system parameters (e.g. kinetic rate constants) and ***s*** are parameters for the readouts, like scaling factors. It can be noted that the parameters ***θ*** = (***p*, *s***) together with the initial conditions ***x***(*t*_0_) and the input ***u***(*t*) fully specify the output from the system.

When experimental data are available corresponding to all or a subset of the system outputs, we denote these data 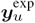, where the index *u* indicates a specific input vector. The corresponding simulated data points from the model (under the same inputs) are denoted 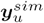. Within this study we only consider steady state output, or output at one specific time point, and therefore from here on we leave out indication of time in the notation. If there are output variables for which we do not have any corresponding experimental data, we denote them 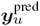.

The problem we would like to address is to describe the uncertainty in the predicted output ***y***^pred^ given the model (1), the data ***y***^exp^, and the prior knowledge we have on the parameters (we drop the index *u* for ease of notation). We would also like to map out those parameters that contribute the most to the prediction uncertainty, as well as those parameters which are important in order to produce qualitatively different predictions (here corresponding to different types of plasticity). To achieve this, we consider the parameters ***θ*** to be stochastic variables (large letters will be used for stochastic variables, e.g. **Θ**)and we useathree step workflow, as illustrated in Figure 1. The workflow consists of (i) inverse uncertainty quantification, (ii) forward uncertainty propagation, and (iii) global sensitivity analysis.

## 2 Background and existing methods

The purpose of **inverse uncertainty quantification** is to estimate unknown parameters of a model from observed data, and at the same time quantify the uncertainty in these parameter estimates. This is most often done in a Bayesian framework (see for example Calderhead and Girolami, 2011; Toni *et al*., 2009; Kramer *et al.,* 2010), by characterizing the posterior distribution, 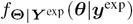, of the parameters. Here ***Y***^exp^ and **Θ** are the stochastic variables corresponding to the experimental data and the parameters, respectively, but for ease of notation we will drop the subscript and refer to the posterior as *f*(***θ**|**y***^exp^). The posterior distribution describes the uncertainty in a set of parameters of a specific model given observed data. The posterior distribution can, by the use of Bayes law, be deduced from the data likelihood *f*(***y***^exp^|***θ***), which describes the likelihood of observing the data ***y***^exp^ from the model given that the parameters ***θ*** are used, and a prior distribution *f*(***θ***), describing the prior knowledge you have about the parameters. The posterior distribution corresponds to

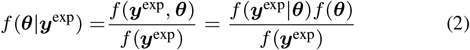

Often the posterior distribution cannot be expressed analytically, rather a sample from the distribution has to be retrieved in order to characterize it. In most cases this is done by the use of Markov chain Monte Carlo (MCMC) methods (Gelman *et al.,* 2013). Furthermore, the standard Bayesian framework is likelihood based, in the sense that we can deduce and compute the data likelihood *f*(***y***^exp^|***θ***). When this is not the case, it is possible to turn to Approximate Bayesian Computation (ABC) (Marjoram *et al.,* 2003; Toni *et al.,* 2009; Sunnåker *et al.,* 2013) which relies on simulation followed by a comparison of simulated and experimental data to assess model fit. In ABC, samples from a prior distribution (or a proposal distribution) are accepted if the experimental data are reproduced by simulations from the model within a certain margin, so that a distance measure *ρ*(*S*(***y***^*sim*^), *S*(***y***^exp^)) is smaller than some predefined cut-off *δ* (*S* is a summary statistic of the data). The accepted parameter sets ***θ*** will form the approximate posterior distribution *f*(***θ**|ρ*(*S*(***y***^exp^), *S*(***y***^*sim*^)) ≤ *δ*). ABC can be used either together with MCMC or with simple rejection sampling. Even when the likelihood expression is known, simulation of the response from a model can be useful to evaluate the likelihood.

The parameter space corresponding to the uncertainty in the parameters is related to what is also referred to as the viable parameter space of a system (Zamora-Sillero *et al.,* 2011), i.e. the subset of the parameter space where a model contains a desirable behaviour. Further approaches to explore the viable space have been described in the literature. In e.g. Gomez-Cabrero *et al*., 2011 particle swarm optimization is used to investigate the viable space.

The extent to which it is possible to deduce values of model parameters via inverse quantification is connected to the identifiability of the parameters. If the true values of the parameters can be deduced from unlimited data, the model is called identifiable. In Raue *et al.,* 2009, identifiability is explored via the so called profile likelihood, and in Vanlier *et al.,* 2012b, the profile likelihood methodology is integrated with a Bayesian approach to deal with non-identifiability.

### Forward uncertainty propagation and global sensitivity analysis

The uncertainty in the model parameters can be propagated to the model predictions, and here be quantified by e.g. the variance of the predictions at a specific time point or steady state. It is of interest to see how this uncertainty in the predictions depends on the uncertainty of specific parameters; i.e. to perform a global sensitivity analysis (GSA) on the predictions based on the posterior distribution. It is not necessarily the case that an uncertain parameter will give uncertain predictions (Gutenkunst *et al*., 2007). In general when performing GSA, the input factors (e.g. model parameters) are assumed to be independent and the GSA is then performed by sampling the factors independently from some marginal distributions (Saltelli *et al*., 2008). The sensitivities are next calculated by e.g. decomposing the output variance based on subgroups of input factors (Sobol, 2001; Saltelli, 2002). Dependencies between parameters make GSA more complex. Methods based on the decomposition of variances can still be used, but the calculation of sensitivities are more expensive and harder to interpret (Saltelli *et al*., 2004). Another approach is to use so called Monte Carlo filtering, in which the output is subdivided into different classes and the respective parameter distributions are compared (Saltelli *et al*., 2004). Other methods, some based on information theory, have also been presented in different studies (Lüdtke *et al*., 2008; Vanlier *et al*., 2012a). More methods for different forms of GSA are reviewed in e.g. Zi, 2011.

## 3 Approach

The approach presented here combines Approximate Bayesian Computation for the inverse uncertainty quantification with decomposition of variance and Monte Carlo filtering for the global sensitivity analysis (Figure 1). We have made some developments to the standard implementations of these methods in order to be able to combine them as well as to make the workflow more efficient, as discussed below.

### Inverse uncertainty quantification through ABC and efficient merging of data

The first step of the workflow consists of characterizing the posterior distribution of the parameters. In order to avoid assumptions of a normal likelihood we use simulation with ABC to sample from the posterior distribution, as non-normal output distributions easily can arise in non-linear systems (Weisse *et al*., 2010).

We use several experimental datasets that are combined in sequence, where the posterior distribution after fitting to one dataset is used as the prior for the fitting to the next, by means of multivariate distributions called copulas (see below and Figure 2). A Markov Chain Monte Carlo (MCMC) approach is used for the ABC sampling (ABC-MCMC) on each dataset in the sequence, and after the final step of the sequence, we also boost the posterior sample by a simple rejection sampling from the final copula. In each ABC-MCMC iteration, we use an adaptive acceptance threshold (or margin) to more efficiently find the viable space where the actual sampling can begin. This is similar to the particle approach proposed by Secrier *et al*., 2009 where the acceptance region is decreased in consecutive runs, although we make this adaption within a single MCMC run.

Copulas are multivariate probability distributions with uniform marginal distributions, which describe the dependence structure between the stochastic variables. Graphical models called vines can be used to formulate copulas that are constructed in pairs in order to describe the dependencies over multiple variables (Bedford and Cooke, 2002) (see also further details in the Supplementary Material). We use D-vines to model the multivariate posterior distributions produced by ABC-MCMC runs. After each step of the fitting sequence described above, a copula is fitted to the posterior sample from that step and is next used as prior for the subsequent dataset. To our knowledge, copulas have not been employed in this way in inverse quantification previously, although they have been used in hybrid proposal distributions in MCMC (Schmidl **et al*.,* 2013). The proposed approach to inverse uncertainty quantification is illustrated in Figure 2.

Based on the posterior distribution, we characterize to what extent different parameters are constrained by the data and model using the sample histograms to assess the entropy of the marginal distributions. The entropy of the sample histogram of the variable *X* was calculated by *H*(*X*) = 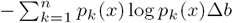, where *p*_*k*_(*x*) is the probability of having the outcome *x* of *X* in the k:th bin, *∆b* is the bin size and *n* the number of bins. The reduction in entropy observed when updating the parameter distributions from the prior to the posterior is used as a measure of the uncertainty decrease of the specific parameters, *H*_diff_ = *H*_prior_ − *H*_post_. The posterior distribution is further characterized by different standard statistical tools like clustered correlation plots and parallel coordinate plots.

### Forward uncertainty propagation and global sensitivity analysis

The next step of the workflow is to translate the uncertainty in the parameters to uncertainty in predictions by performing simulations based on all parameter sets in the posterior distribution sample. The uncertainty of the predictions ***Y***^pred^ is next quantified by the variance of each vector element *V*(*Y*^pred^).

Finally, we perform a global sensitivity analysis to investigate from where the uncertainty in the prediction stems. This is done in two ways with two different aims. First, we investigate which parameters on average reduce the uncertainty in the prediction the most if they were known more precisely. Second, we look into which parameters are most influential in separating different qualitative behaviours of the model.

In order to address the first aim we decompose the variance of the output based on the contribution from different input factors of the model (model parameters in our case). The first order sensitivity index (Saltelli *et al*., 2004) quantifies the impact that a model parameter has on a specific output, and is defined by 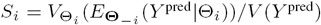. Here ***Θ***_–*i*_ stands for all parameters of the vector **Θ**, except *Θ*_*i*_. The expression 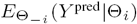 is thus the expected value of *Y*^pred^ over all parameters except *Θ*_*i*_ when *Θ*_*i*_ is conditioned on a specific value 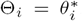, and 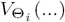 is the variance over all specific values 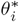. Well established and efficient methods to calculate *S_i_* for distributions of independent input factors (Sobol, 2001; Saltelli, 2002) are available. However, since we are performing the GSA using the multivariate posterior distribution, *f*(***θ***|***y***^exp^), which displays dependencies between possible model parameter values due to the inner structure of the model, these methods cannot readily be applied. Instead, we perform a calculation inspired by (Saltelli *et al*. (2004), chapter 5.10), but with modifications in order to utilize the already existing posterior sample produced from the ABC method. This computation is based on binning the posterior space and results in approximation of the sensitivity index *S*_*i*_ (details can be found in the Supplementary Material).

If the model is not sufficiently constrained by the experimental data, a large variance can be seen in the prediction and qualitatively different output behaviours can be observed. It is then of interest to identify the parameters with the largest impact in separating these behaviors. This is known as Monte Carlo filtering (Saltelli *et al*., 2004). In order to do this we first group the predictions into classes with different qualitative behaviour, and also divide the posterior distribution sample according to the same grouping. Model parameters that have a large influence on the model behavior in question display different sample distributions in the different groups. We consider marginal as well as pairwise parameter distributions, and sort them based on the Kolmogorov-Smirnov test and Kullback-Leibler divergence, respectively.

**Fig. 2.**
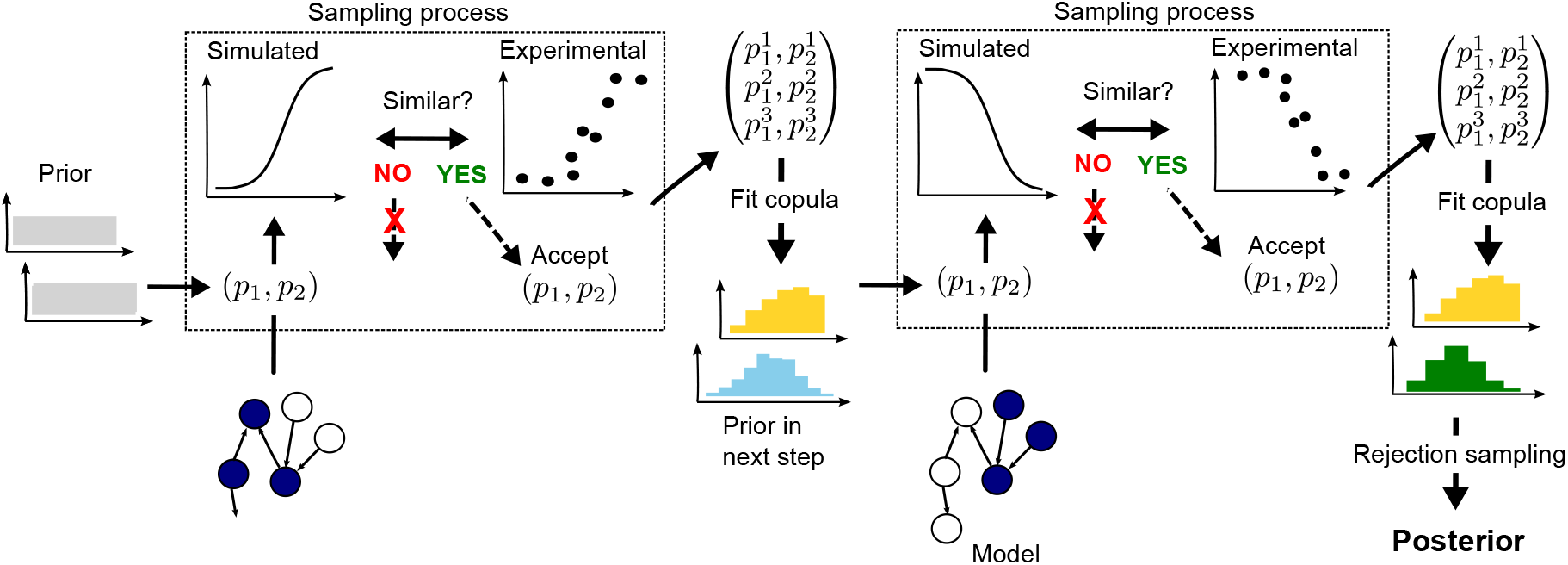
Sequential approach for mapping out the viable space when multiple experimental datasets corresponding to different experimental setups are available. In this case we have two experimental setups available (used in each of the dashed boxes and exemplified by different parts of the model being active as highlighted in blue), and exemplify the approach for two parameters. The model calibration is done in steps, so that we start with a uniform prior for the first dataset, which produces a posterior distribution used as the prior in the next step by fitting and sampling from a multivariate copula model of the distribution. The process shown within the dashed boxes (draw from prior, simulate data, compare to experimental data and keep/not keep parameter set) is repeated many times using MCMC. The posterior distribution from the sequential fitting with two phenotypes is produced after the final MCMC step by rejection sampling from a copula fitted from the distribution produced by MCMC.

## 4 Application

We have applied our approach to a previously constructed intracellular model that in a simplified way exemplifies a molecular mechanism important for the strengthening (long term potentiation, LTP) or weakening (long term depression, LTD) of neuronal synapses (Nair *et al*., 2014). The modification of synapses through the process of LTP or LTD is a complicated process including a number of kinases, phosphatases and scaffolding proteins (Woolfrey and Dell’Acqua, 2015). This process is, however, often assumed to be effectuated by the balance between a few important kinase and phosphatase enzymes, and in the model used in this study (Nair *et al*., 2014), this balance is due to the interaction between calcium (Ca), calmodulin (CaM), which contains four Ca-binding domains, protein phosphatase 2B (PP2B, also known as Calcineurin), Ca/CaM-dependent protein kinase II (CaMKII) and protein phosphatase 1 (PP1), as illustrated in Figure 3.

### 4.1 Model

The model consists of 25 species (corresponding to proteins, protein complexes, the activated form of a protein or the input signal Calcium) and 34 reactions, were all reactions except two are elementary reversible reactions based on the law of mass action. This means that the reactions are of the type: *A* + *B* ⇄ *C,* where *A, B* and *C* are different species, where the right going reaction has a kinetic constant denoted *k*_*f*_ and the reaction in the opposite direction has a kinetic constant denoted *k*_*r*_. We also use the equilibrium constants *K*_*d*_ = *k*_*r*_/*k*_*f*_. All species and reactions are listed in Table S1 and Table S2, respectively. There are also thermodynamical constraints which apply when there is more than one reaction path between a pair of species. These are expressed by the so called Wegscheider conditions (Wegscheider, 1911; Gorban and Yablonsky, 2011; Yablonskii, 1991) and link some *K*_*d*_, parameters of the model to other *K*_*d*_, parameters (Table S3). We therefore decompose the *K*_*d*_-parameters in the model into two sets; free *K*_*d*_-parameters that are modified throughout the analysis, and thermo-constrained *K*_*d*_-parameters whose values are set by the values of the free parameters via these rules. More information about the model can be found in Section S1 in the Supplementary Material.

**Fig. 3.**
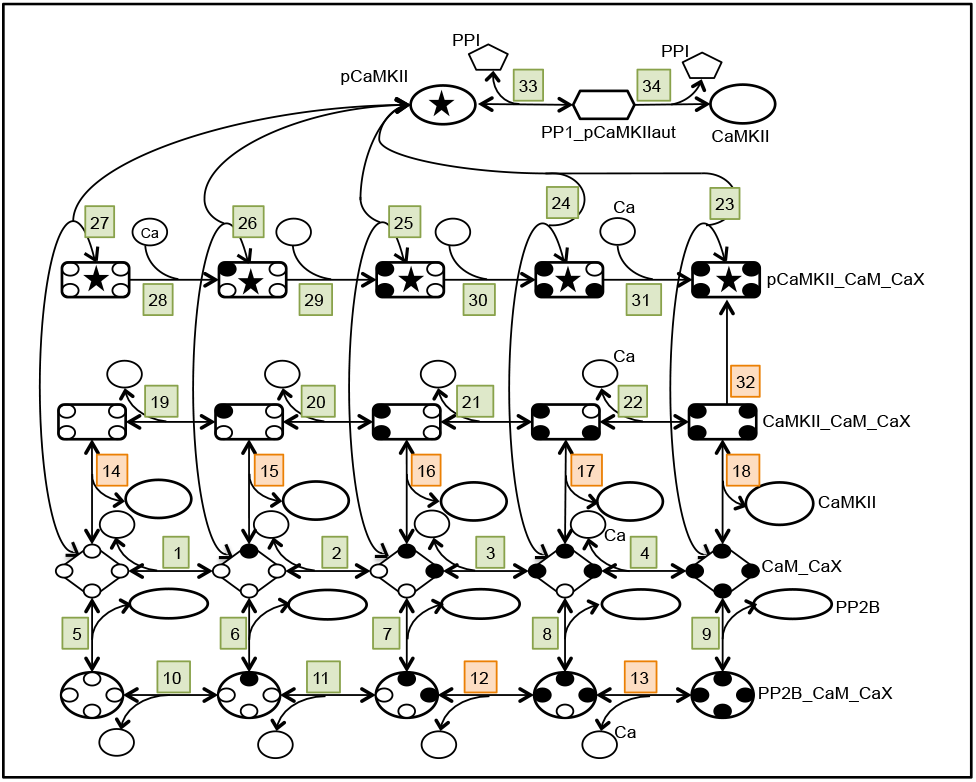
A graphical representation of the intracellular model used to illustrate the proposed approach to model characterization. The numbers indicate the corresponding reactions in Table S2. The small black filled circles correspond to Ca domains that have bound a Ca and the X in CaX correspondingly denotes Ca,…,Ca4 (CaO is simply referred to as Ca in Table S2). Reactions colored in orange correspond to parameters that would be the best targets to identify in order to reduce prediction uncertainty, and correspond to the parameters in the legend of Figure 6.

### 4.2 Experimental data for parameter estimation

The parameter estimation was based on quantitative data collected from a number of publications as described in Nair *et al*., 2014. The data correspond to different experimental setups describing different, experimentally engineered, phenotypes of the system. The phenotypes correspond to a subpart of the system and are characterized by steady state (or close to steady state) input-output curves, where, in each experiment, a given input is varied in value in order to obtain the curve. In the model, the experimental phenotypes are recreated by applying different model inputs ***u**.* The different phenotypes, and the subparts of the model that are active under the different settings, are described in Table S2.

### 4.3 Prior distributions

We obtained default values for the free parameters from Nair *et al*., 2014 and used them as the centers *µ*_*i*_ for log-uniform prior distributions. The range of the prior was set as *µ*_*i*_ – 3 to *µ*_*i*_ + 3 in log-space. We did not sample the thermodynamically constrained parameters, nevertheless, we can assign implicit prior distributions to them via the thermodynamic constraint rules. By sampling the free parameters and propagating these through the rules, we can obtain prior samples for the constrained parameters as is shown in Figure S1.

### 4.4 Model reduction

The different phenotypes correspond to situations either very close to steady state or with slow dynamics, and we have utilized this fact in order to reduce the model. For some of the phenotypes (phenotypes 1-4 in Table S2) the output could be approximated with steady state. Steady state reduction was hence performed to the model in order to speed up calculations, resulting in analytical steady state solutions for subparts of the model. The reduction was based on the principle of detailed balance (Yablonskii, 1991), which has the consequence that steady state only can occur at an equilibrium and so all reaction fluxes are zero. Since the reaction fluxes are of the form *k*_*f*_[*A*][*B*] – *k*_*r*_ [*C*], it follows that the equilibrium concentrations of the species depend only on *k*_*r*_/*k*_*f*_ = *K*_*d*_ (using that the equilibrium equations can be rewritten as log[*A*] + log[*B*] – log[*C*] = log(*k*_*r*_/*k*_*f*_) = log(*K*_*d*_)). In this way, the equilibrium equations were solved analytically, while making use of the mass conservation laws of the system (i.e. that the total amount of each elementary species remains the same during the experiment). This enabled us to express the equilibrium concentrations as functions of the *K*_*d*_ parameters and the total amounts of the species. More information about the analytical solutions can be found in the Supplementary Material.

For the remaining phenotypes (phenotypes 5-6 in Table S2) we utilized the fact that they have slow dynamics, and that the output therefore mainly should depend on the *K*_*d*_ parameters. The problem was thereby reduced to first finding the posterior for the *K*_*d*_ parameters, based on constant *K*_*f*_, and then expand this posterior to *k*_*f*_:s by simple rejection sampling.

### 4.5 Results

#### Inverse uncertainty quantification and characterization of the viable space

Given the prior distributions, experimental data and model structure, a sample from the posterior distribution was retrieved through the sequential ABC-method that had a good fit to the experimental data (details on the distance measure and normalization procedures used can be found in the Supplementary Material). The sampling was performed on a parameter log-scale and the multivariate posterior distribution was characterized by looking at single parameters as well as pairs of parameters.

The marginal posterior distributions of all *K*_*d*_ parameters are summarized in the parallel coordinate plot of Figure 4, where the prior distribution and reduction in entropy also are indicated. The forward *k*_*f*_ and backward *k*_*r*_ parameters are not included in the figure since these had, as expected since we use mainly steady state data to fit the model, a posterior distribution very similar to the prior (and a corresponding low reduction in entropy). It can be noted that some parameters are very constrained by the model and currently used data, with the two most prominent examples being K_d_*CaM_Ca4*PP2B and K_d_*CaM*PP2B, which both have a narrow distribution and a large reduction in entropy. Other parameters instead occupy the parameter space up to the edge of the prior, e.g. kautMax and K_d_*CaMKII_CaM_Ca1*Ca. This could be a sign of the prior being too small to include the full viable space or a sign of non-identifiability which is (artificially) resolved by imposing a prior.

**Fig. 4.**
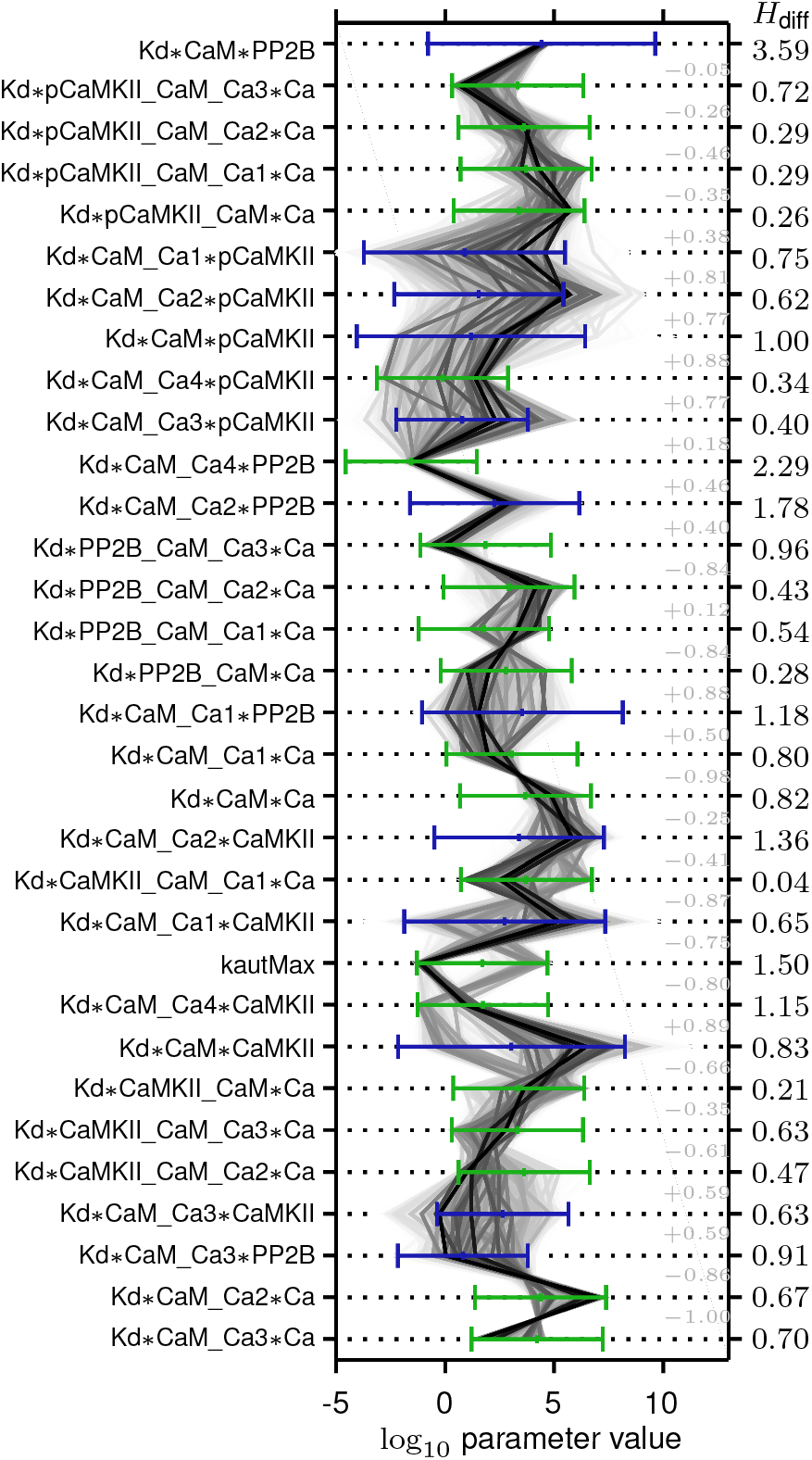
Illustration of the marginal posterior distribution and reduction in entropy for all parameters. The numbers indicated to the far right correspond to the reduction in entropy (*H*_diff_) when going from prior to posterior distribution, and the light grey numbers correspond to the pairwise correlations. Each sample in the posterior distribution is connected across the parameters by a thin grey line, the darkness of which reflects to the posterior probability density at that sample point (the kernel density estimate). The prior of the free parameters is indicated by green bars (showing the range of the log-uniform distribution) and the prior of the thermo-constrained parameters (calculated through the equations of Table S3) is indicated by blue bars (showing one standard deviation of the, lognormal-looking, distributions).

Most parameters that have a narrow posterior distribution (like K_d_*CaM_Ca4*PP2B) display a corresponding large reduction in entropy and vice versa. It can however be noted that some parameters have a wide posterior distribution even though they at the same time have a large reduction in entropy (kautMax and K _d_*CaM_Ca4*CaMKII) which is due to a prominent bimodality in the marginal distributions (Figures 4 and S1). Bimodal distributions can contain a lot of information about the parameter, despite possibly having a wide spread. Histograms of the marginal posterior distributions and their characteristics, such as model-based credibility intervals, are given in Figure S1 and Tables S4-S5.

We also examined possible couplings between parameters by a clustered correlation plot (Figure 5), where the parameters are clustered into groups based on their correlation profile. Some parameter pairs show large correlations, while most others appear to be uncorrelated or only weakly correlated. The pattern of correlations between the parameters of the model also have a tendency to follow the model structure, so that parameters with a high value in the correlation plot are close in the graph of Figure 3. Interestingly, the correlations involving the bimodal parameters are clustered into two separate groups indicated by blue frames in Figure 5, and we observe that the bimodality pattern likely originates from at least two sources in the model since there are no correlations between parameters of one of these clusters to the other.

**Fig. 5.**
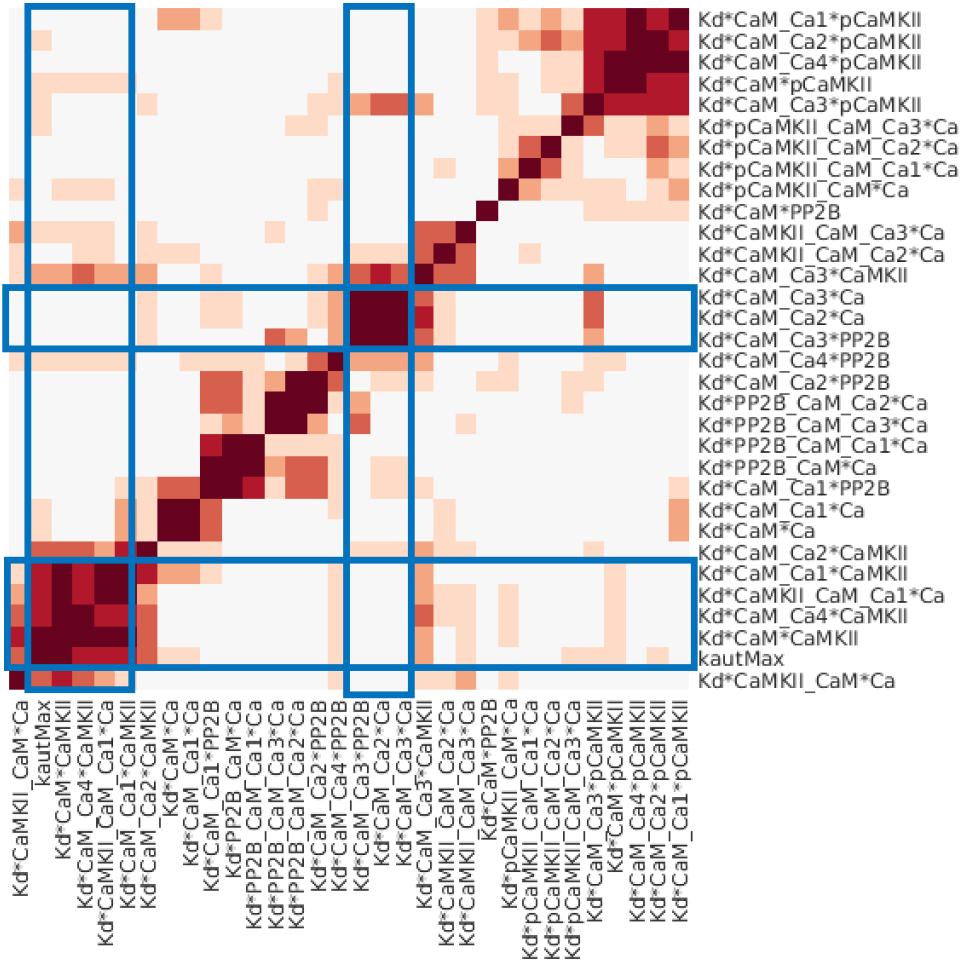
Correlation plot of model parameters based on the samples from the posterior distribution. The parameters are clustered based on their correlation profile, i.e. the array of correlation coefficients for each parameter, via hierarchical clustering with an Euclidean distance metric and average linkage. The blue lines frame the arrays with correlations related to the bimodal parameters.

**Fig. 6.**
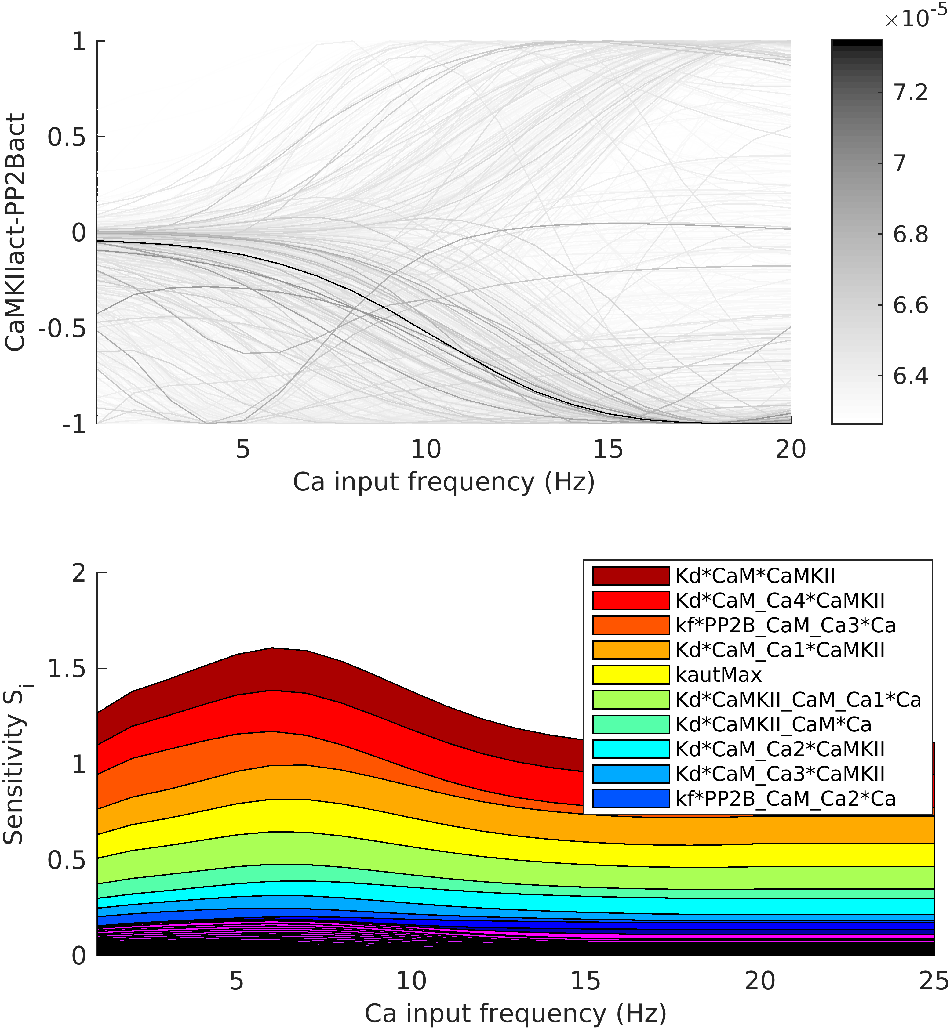
Uncertainty and sensitivity of the prediction. The prediction corresponds to the (normalized) difference between the activity of CaMKII and PP2B at different Ca frequencies. Top panel: The different outputs (grey lines) correspond to different sample points from the posterior distribution. A large uncertainty in the prediction can be observed. Bottom panel: First order sensitivity index (*S*_*i*_) for all parameters at different input (Ca frequency) values. Some parameters have a large influence on the uncertainty.

#### Forward uncertainty propagation

We next analysed how the uncertainty in the parameters is propagated to uncertainty in the prediction that we would like to make from the model. The prediction used here to demonstrate the workflow corresponds to the relationship between the active form of the kinase CaMKII and the active form of the phosphatase PP2B and how this relationship depends on the frequency of Ca transients given as input (for details on input and output functions see supplementary material). The presence of a large amount of activated CaMKII relative to activated PP2B will give long term potentiation (LTP) with the reverse relationship instead resulting in long term depression (LTD).

For all parameter sets of the posterior distribution we calculated the corresponding CaMKII-PP2B relationship at different Ca frequencies (Figure 6). There is a large variation in the prediction given a certain Ca frequency, showing that the model, with currently used data, is not sufficiently constrained to give a precise prediction. In order to investigate the best way to reduce this uncertainty and to learn more about the system we next performed a global sensitivity analysis.

#### Global sensitivity analysis

First we analysed how the uncertainty in the different parameters contributes to the uncertainty of the prediction by decomposing the variance of the prediction based on the different single parameters (see Approach). The single parameters that, if known, on average would give the largest reduction in the uncertainty of the prediction are shown in the legend of Figure 6 and are also indicated in Figure 3. The dissociation constants corresponding to the binding of CaM to CaMKII (reactions 14-18 in Figure 3 and Table S2), as well as the CamKII-CaM complex binding the first two Ca (reactions 19 and 20) are important. Also the maximal autophosphorylation rate (reaction 32), and CaM bound to PP2B binding the third and fourth Ca (reaction 12 and 13) would give a large reduction in uncertainty. It can be noted that several of these parameters are correlated within the posterior distribution (Figure 5). This means that knowledge about one parameter would automatically decrease the uncertainty in the other correlated parameters as well. One possible approach for further investigation would therefore be to identify the most sensitive parameter of each cluster of correlated parameters and try to experimentally determine its value. The remaining parameters in each cluster would then likely also show a large decrease in uncertainty.

The next part of the GSA was to analyze qualitatively different types of output behaviors via Monte Carlo filtering. We show the output corresponding to the prediction as well as the output for phenotype 5 (Table S2) in Figures 7-8. Phenotype 5 was included since it exhibits a somewhat peculiar behavior. Both outputs were divided into two different classes (Figures 7-8, top panels) depending on whether, in the case of phenotype 5, the output was monotonic or not, and for the prediction, whether or not the output agreed with a hypothesized behaviour for synaptic plasticity. The sample from the posterior distribution was also subdivided according to the same classes and analysed both at the individual parameter level and by investigating all parameter pairs. (Figures 7-8, bottom panels). The individual parameters as well as the parameter pairs were sorted based on the distance between the distributions when comparing the two classes. A Kolmogorov-Smirnov test was used for the marginal distributions for the individual parameters, and the Kullback-Leibler divergence was employed for the joint distribution of parameter pairs.

For phenotype 5, an interesting example is the parameter K_d_*CaM*CaMKII (Figure 7, bottom left), for small values of this parameter only monotonic output curves are observed, but for larger values we have both types of curves. When pairs of parameters are considered, the scatterplot corresponding to the pair with the largest KLD distance, K_d_*CaM*pCaMKII versus K_d_*CaM_Ca4*pCaMKII, shows an interesting separation between classes, not visible in the marginal histograms (Figure 7, bottom right). Further histograms and two dimensional projections of top scoring parameters and pairs are shown in Figure S2 and Figure S4.

**Fig. 7.**
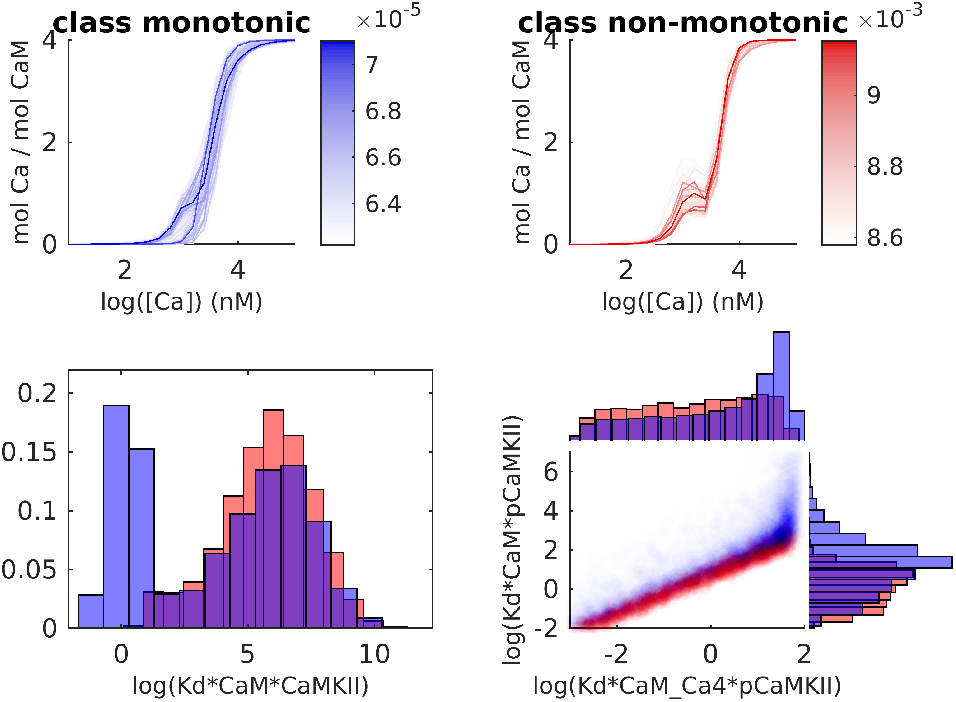
Classification of outputs from phenotype 5 and corresponding subdivision of the posterior distribution. Red corresponds to non-monotonous output while blue corresponds to the monotonous output curves. Top panel: Subdivision of the output mol Ca per mol CaM into a monotonous and non monotonous group. Bottom left panel: Marginal histograms for a parameter with a large difference in the posterior distribution between the two classes, as quantified by the Kolmogorov-Smirnov test. Bottom right: Pairwise scatterplots of of the parameter pair with the largest KLD-distance.

**Fig. 8.**
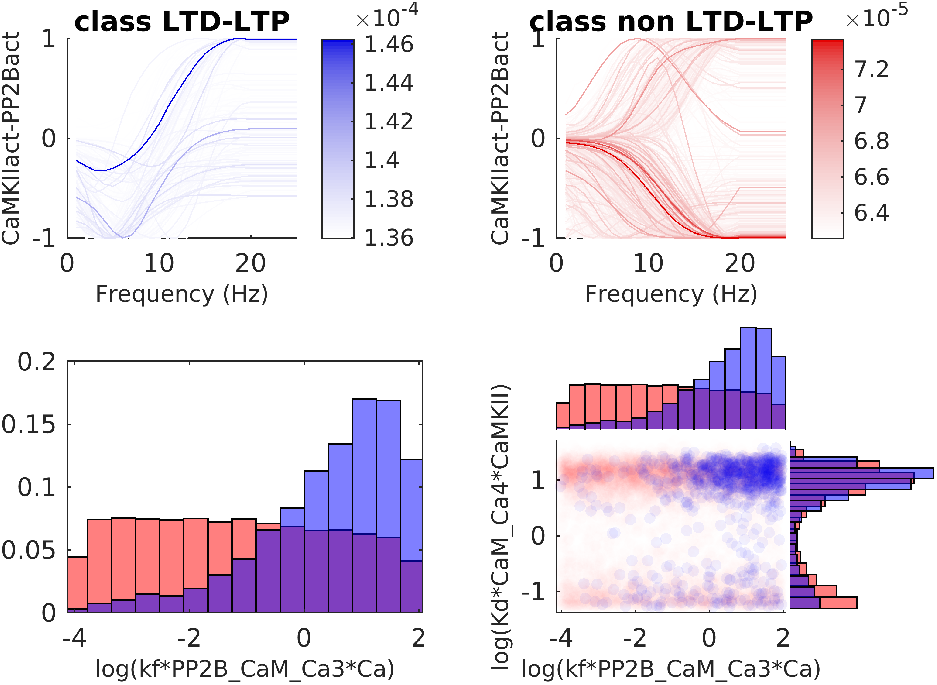
Classification of predictions and corresponding subdivision of the posterior distribution. Blue corresponds to the hypothesized output behaviour for synaptic plasticity (LTD-LTP) while red corresponds to the curves that do not follow the hypothesized behaviour (non LTD-LTP). Top panel: Subdivision of the prediction output according to the two classes. Bottom left panel: Marginal histograms for the the parameter with the largest difference in posterior distribution between the two classes quantified by a Kolmogorov-Smirnov test. Bottom right panel: Pairwise scatterplots for the highest KLD-scoring pair.

For the prediction, the hypothesized output behaviour for synaptic plasticity describes a specific relationship between Ca level and kinase-phosphatase activity (denoted CamKIIact-PP2Bact, in Figure 8).This relationship is as follows. At very low levels of Ca there is a balanced kinase-phosphatase activity: for low levels there is a negative balance (i.e. kinases dominate), and for larger levels of Ca there is a positive balance (i.e. phosphatases dominate). The classification criterion, which was based on whether this behavior is shown or not, is formalized mathematically in Section S3.3 in the Supplementary Material.

Similarly, for this output we observe differences between the two classes with respect to the marginal parameter distributions (see Figure S3 for histograms of top scoring parameters), but for this output, the number of parameters with a clear separation between the two class distributions are much fewer than in the earlier case with phenotype 5. The bottom left panel of Figure 8 shows the histogram of the parameter with the largest distance between the classes. For k_f_*PP2B_CaM_Ca3*Ca there is a larger probability of receiving the hypothesized behavior if k_f_*PP2B_CaM_Ca3*Ca> 10^-1^. Looking at the top scoring KLD-pairs (Figure S6) does not provide any extra information as compared to the histograms. The highest scoring pair is shown in the bottom right panel of Figure 8.

## 5 Discussion

We have here presented a workflow for analysing the viable space of biochemical models, using a previously constructed model of CaMKII and PP2B as an example. By combining Bayesian analysis with global sensitivity analysis we can quantify the uncertainty in the model parameter estimates and model predictions, as well as pinpoint where this uncertainty stems from. This is useful both for experimental design as well as model building. This is also interesting from a sensitivity analysis perspective.

Biochemical models are generally uncertain (Geris *et al*., 2016) with a large viable space. Performing an a-priori sensitivity analysis, e.g. based on intervals, seems not feasible for a model of this size. Similarly sensitivity analysis of a product space of posterior intervals would lead to model behavior far outside the bounds set by the data and lead to errors. When GSA is performed within the full posterior distribution it takes the correlations between parameters into account and only investigates data fitting parameters.

Analysing the viable space of complex models with many parameters is, however, computationally expensive. By the use of model reduction as well as integrating data sets sequentially with copulas we could reduce the computational cost to the point where an extensive analysis could be performed.

A Bayesian approach together with GSA is of course more rigorous than a manual parameter search, since it accounts for the variability in parameter space. It thereby provides more accurate and extensive predictions. It also offers predictions on parameter regions (e.g. levels of kinetic constants) which are correlated with desired behaviours. An additional value is that a the prior distribution makes the assumptions on modeling uncertainty more explicit, which is more useful when sharing and comparing models than a single, seemingly working parameterization.

There are other workflows of model analysis described in the litterature which e.g. focus on fast optimization and statistical classification and clustering techniques (Gomez-Cabrero *et al*. (2011)), while the procedures presented here aim to handle uncertainty quantification and propagation consistently through all steps. On the other hand, several similar, statistically embedded experiment design methods (e.g. Liepe *et al*. (2013); Weber *et al*. (2012)) focus on maximizing information in planned experiments, whereas this analysis workflow assigns roles to model constituents and measures of importance to parameters. Ultimately we attempt to better understand the mechanisms in the model. This understanding can of course be used for experiment design, so these topics are highly connected.

We have so far only spoken about the viable space in terms of model uncertainty due to missing data. Another reason that a viable space is a better description than a single parameter vector is biological variability because biological measurement techniques often target cell populations rather than single cells. Biochemical pathway models, on the other hand, often correspond to a generic individual cell or cell compartment. With a Bayesian approach it is possible to capture (smoothly) varying biological properties, even though it cannot distinguish between uncertainty due to missing data and biological variability.

## Acknowledgements

The simulations were performed on resources provided by the Swedish National Infrastructure for Computing (SNIC) at Lunarc, and we thank Anders Sjöström at Lunarc for valuable technical support.

## Supplement

### S1 Model details

#### S1.1 Additional information on the LTP LTD pathway - autophosphorylation

As described in the main text, the following elementary species are included in the model: calcium (Ca), calmodulin (CaM), protein phosphatase 2B (PP2B) and Ca/CaM-dependent protein kinase II (CaMKII) and protein phosphatase 1 (PP1). CaM is a Ca-binding protein involved in multiple signaling processes and is strongly implicated in synaptic plasticity. CaM contains four Ca-binding domains, each binding one Ca ion. The binding of Ca by CaM is a cooperative process. Ca-bound CaM activates PP2B, another protein implicated in molecular processes related to learning which also plays a role in striatal signaling. The third protein, CaMKII, is a kinase, which is activated by the binding of Ca-CaM. CaMKII molecules exist as dodecamers, consisting of two hexamer rings. A CaMKII unit that has bound CaM can autophosphorylate when sitting beside an active neighbouring unit in the same hexamer ring. The phosphorylated unit can remain active even in the absence of Ca-CaM.

#### S1.2 Additional information about the model - reaction 32 (autophosphorylation) and reaction 34

Two of the model reactions are not elementary reversible reactions of the type described in the main text; one of these is different only in the way that it is irreversible (reaction 34 in Table S2), the other one is more complicated as it describes the autophosphorylation process of CaMKII monomers.This process is for practical reasons reduced with the help of a phenomenological rate function, and corresponds to reaction 32 in Table S2. The rate function *f*_*aut*_ (*x*) = *k*_*autmax*_*g*_*aut*_(*x*) describes how much active CaMKII units that are autophosphorylated each time step as a function of the proportion of active CaMKII monomers, *x.* It consists of a constant *k*_*autmax*_ corresponding to the maximum rate, times a function

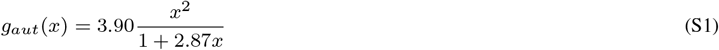

describing the probability that an activated CaMKII monomer has another activated monomer as a neighbour (Li *et al*., 2012). The constant values within the function *g*_*aut*_ were retrieved by fitting to the data of Figure S3 in Li *et al*., 2012.

##### S1.2.1 Input *u*(*t*), and output functions *y*(*t*) = *g*(*x*(*t*), *s*)

The input functions correspond to ***u***(***t***) = (Ca(*t*), CaM(*t*)). For each experimental/in silico condition only one of the inputs are varied and the other held constant, as described in Table S2. For the first six conditions (phenotypes 1-6) the inputs are also constant in time, for the last condition (the prediction) the input, *u*_*Ca*_(*t*), is a sequence of 10 spikes (a so called spiketrain), with frequency, f, starting at *t* = 0 (Nair *et al*., 2014) (for detailed description on the spiketrain, see Nair *et al*., 2014 Figure 12.3 and Figure 12.4).

Let ***x***(*t*) be a vector corresponding of all model species (state variables) stated in Table S1, and define:

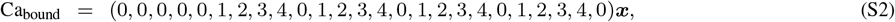

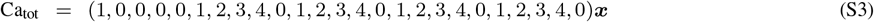

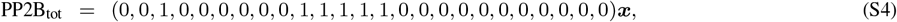

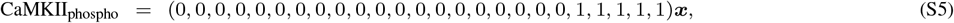

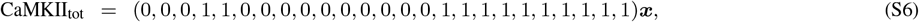

and

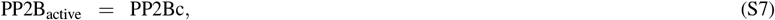

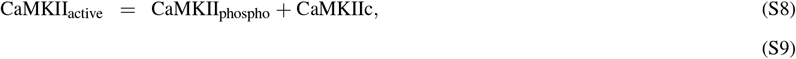

then the outputs of the different experimental settings correspond to

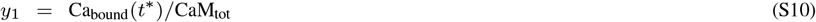

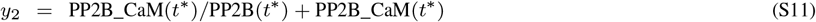

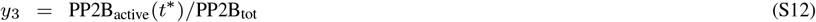

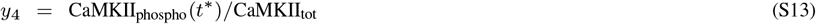

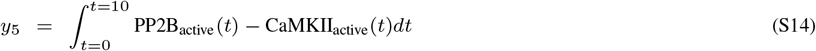

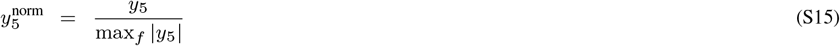

where *t\** in our simulations were set to *t* =* 600, which we assumed where near steady state, and *f* is the frequency of the spike train *u*_*Ca*_(*t*).

**Table S1.** The different substances in the system presented by Nair et al., 2014.

| Number | Name |
| --- | --- |
| 1 | Ca |
| 2 | CaM |
| 3 | PP2B |
| 4 | CaMKII |
| 5 | pCaMKII |
| 6 | CaM_Ca1 |
| 7 | CaM_Ca2 |
| 8 | CaM_Ca3 |
| 9 | CaM_Ca4 |
| 10 | PP2B_CaM |
| 11 | PP2B_CaM_Ca1 |
| 12 | PP2B_CaM_Ca2 |
| 13 | PP2B_CaM_Ca3 |
| 14 | PP2B_CaM_Ca4 |
| 15 | CaMKII_CaM |
| 16 | CaMKII_CaM_Ca1 |
| 17 | CaMKII_CaM_Ca2 |
| 18 | CaMKII_CaM_Ca3 |
| 19 | CaMKII_CaM_Ca4 |
| 20 | pCaMKII_CaM |
| 21 | pCaMKII_CaM_Ca1 |
| 22 | pCaMKII_CaM_Ca2 |
| 23 | pCaMKII_CaM_Ca3 |
| 24 | pCaMKII_CaM_Ca4 |
| 25 | PP1 |

**Table S2.**
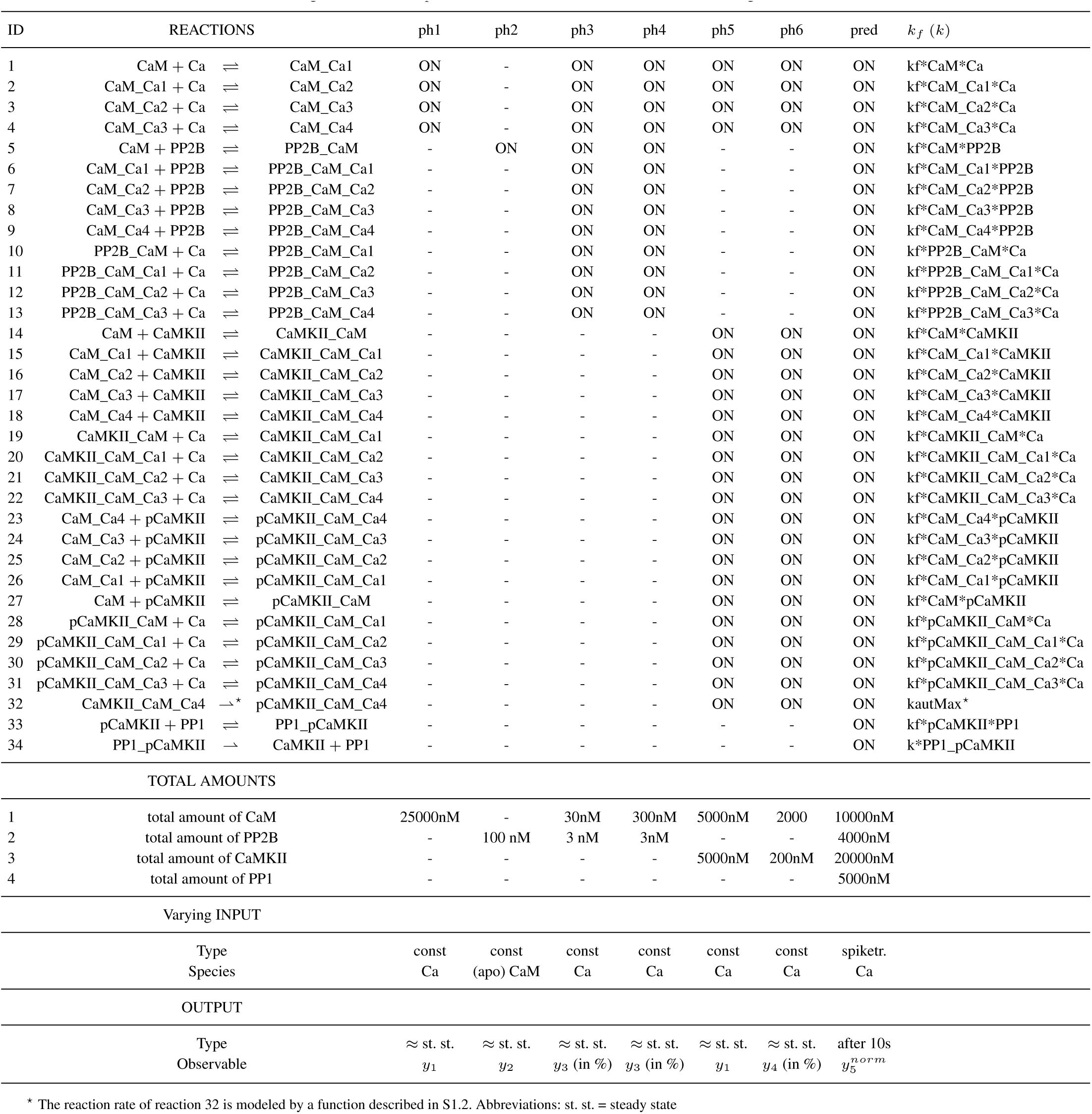
Summary of chemical reactions, total amounts, inputs and outputs of the system described in Nair et al., 2014. The different observed phenotypes (denoted ph1-ph6) correspond to different experimental set ups. The prediction (pred) corresponds to Figure 12.4C of Nair et al., 2014. All reactions have reaction rates based on the law of mass action, except the one marked with * where the reaction is more complex (see main text for details). A list of all species can be found in Table S1. By approximately steady state (≈ st. st.) we mean an experiment/simulation with a long duration (several minutes). The names of the forward reaction kinetic constants *k*_*f*_ are given, the names for *k*_*r*_ and *K*_*d*_ for the reversible reactions follow the same naming convention. The parameters of the two irreversible reactions are also given.

**Table S3.**
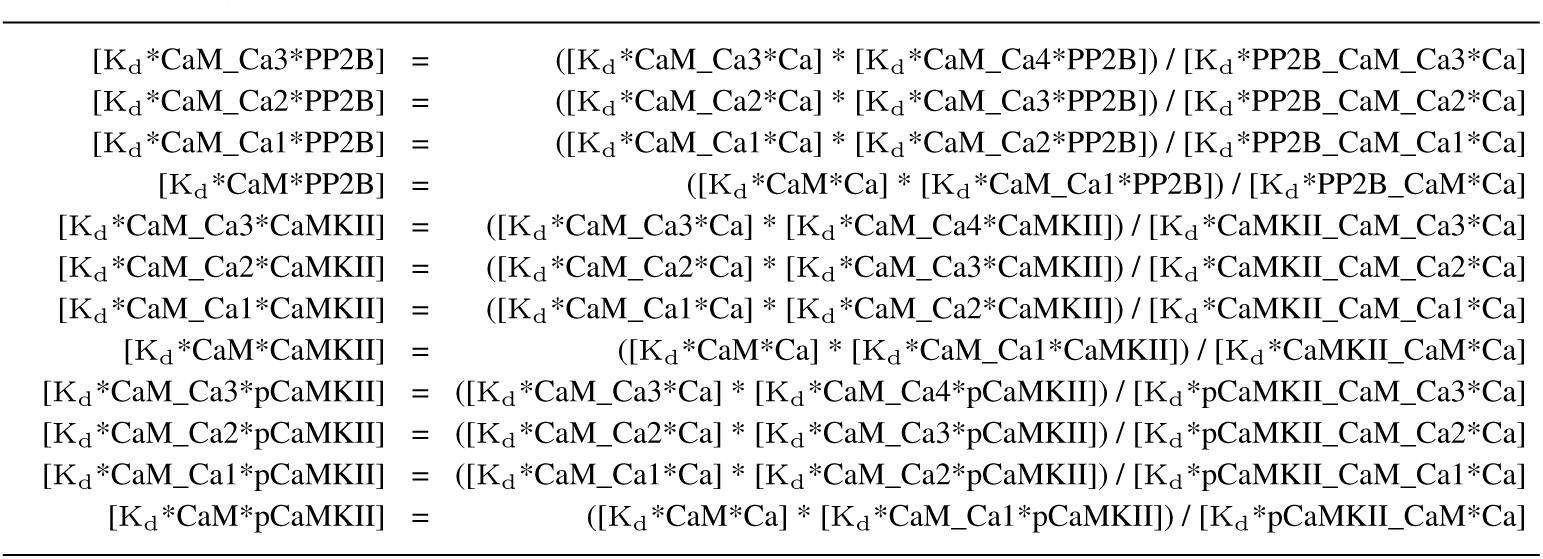
The thermodynamic constraint rules connecting different equilibrium constants of the model due to multiple possible reaction paths from one species to be converted to another.

### S2 Details of the ABC-MCMC uncertainty quantification

#### S2.1 A short introduction to copulas

We are interested in describing the multivariate distribution for the random vector ***X*** = (*X*_1_,…, *X*_*d*_)’ in some way. The elements of ***X*** have continuous marginal distribution functions; *F*_*i*_(*x*_*i*_) = *P*(*X*_*i*_ ≤ *x*_*i*_). A copula is a function that connects the multivariate distribution function to the marginal ones (*F*_*i*_) as follows.

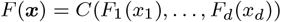

It can be shown (Sklar’s theorem) that for continuous marginal distributions, *C* is unique. The elements of the vector (*U*_1_,…, *U*_*d*_) = (*F*_1_ (*x*_1_),…, *F*_*d*_(*x*_*d*_)) are by definition uniformly distributed. Hence copulas can be viewed as multivariate distribution functions whose one-dimensional margins are uniform on the interval [0, 1] (Nelsen, 2006).

The pair-copula decomposition of a multivariate distribution is useful in order to describe the distribution in question. Consider the vector ***X*** and the corresponding density function *f*(***x***) = *f*(*x*_1_,…, *x*_*d*_) which can be factorized as

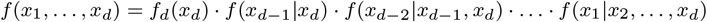

It can be shown that the multivariate (joint) density can be represented by a number of appropriate pair-copulas times the conditional marginal densities based on this factorization. For a vector with three components, we have for example (by use of the chain rule)

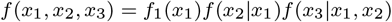

and

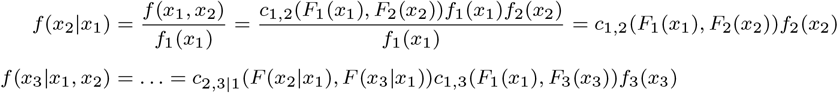

This gives

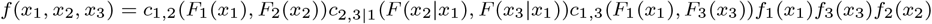

The copula pairs can be chosen independently of each other giving a wide range of possible dependence structures, especially for high-dimensional distributions. Graphical models called vines were introduced to arrange the pair copulas in a tree structure (see e.g Bedford and Cooke, 2002 and Aas *et al*., 2009). C- and D-vines are constructed by choosing a specific order of the variables included.

#### S2.2 Normalization

In order to be able to compare different experimental setups, as well as normalizing on a log scale for the input, the input *x*_*sim,j*_ and output *y*_*sim,j*_ in each simulation was normalized according to

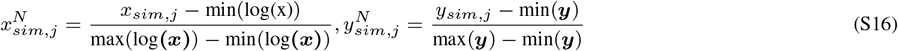

where 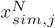 is the normalized version of the ith component *x*_*sim,j*_, and similar for 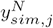. The experimental data ***y***_exp_ and the experimental input (on the log-scale) ***x***_exp_ were then normalized with the same quantities as for the simulated data.

#### S2.3 Distance measure

We have used the following distance measure:

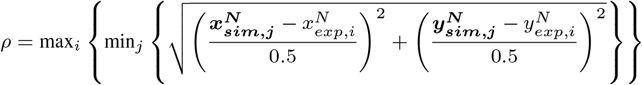

where 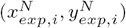 is the normalized experimental data point *i, i =* 1…*n* and 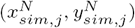 is the normalized simulated data point *j, j* = 1…m, and *m* >> *n.* The simulated data points was retrieved using a dense grid on the x-axis (in the order of 1000 points). Checking whether *ρ < δ,* where *δ* is the chosen ABC-threshold, corresponds to defining a circles around each normalized experimental point and checking that all circles has a part of the simulated curve passing through. The circles has a radius equal to 0.5*δ*, which is a deviation of 100*δ*% of the average normalized output (0.5) for all points on the curve, The value of *δ* was set to 0.1, corresponding to a 10% deviation. We adopt this scheme in order to account for noise in both input and output variables.

### S3 Global sensitivity analysis

#### S3.1 Normalisation and scale

The output from the prediction, *y_7_*, which can be both positive and negative, were normalised according to 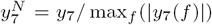, where *f* is the input frequency of Ca-spikes (for a detailed description of the input, see Nair *et al*., 2014. The binning in the sensitivity analysis was performed on the log10-scale.

#### S3.2 Sensitivity indices

We here describe the calculation of the sensitivity index *S*_*i*_, of the parameter Θ_i_ and the scalar output, *Y,* over the posterior distribution 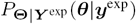, where **Θ** is a vector corresponding to the model parameters, and ***Y***^exp^ a vector corresponding to the output variables that vi have experimental data from. The sensitivity index is defined by 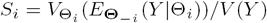, where ***Θ***_–*i*_ corresponds to all elements of ***Θ*** except *Θ*_*i*_.

This calculation was inspired by Saltelli *et al*., 2004 (chapter 5.10), but with major modifications in order to use the existing posterior sample produced from the ABC. The prediction is a (complicated) function of the parameters *y = h(**θ**)),* i.e. each sample point ***θ*** has a corresponding ***y*** value (*y* also depend on the input ***u,*** but to simplify the argument we here assume a specific input, ***u*** = ***u**\**). We first want to calculate the inner conditional expected value 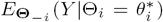, in principle for all different values of 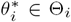. We approximate this by binning the posterior sample with respect to *Θ*_*i*_, into *m* number of bins of size *σ,* with midpoint 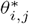, and indexed by 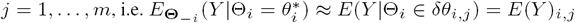, where 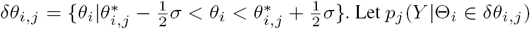 be the conditional distribution of *Y* given that *Θ*_*i*_ is in the *j*:th bin. We denote the corresponding subsample from the posterior distribution 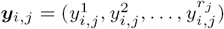, where *r*_*j*_ is the total number of points in the j:th bin. The sample conditional and unconditional means are:

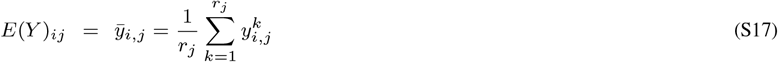

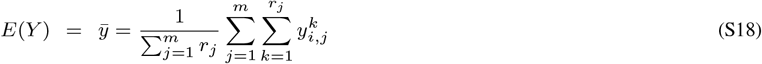

The sensitivity index is calculated by:

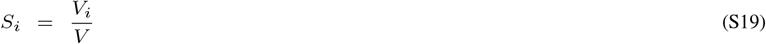

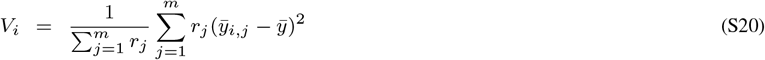

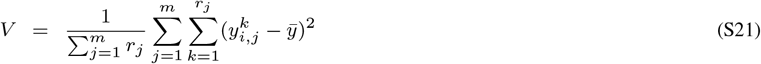

#### S3.3 Monte Carlo filtering

##### S3.3.1 Classification of the prediction output

The outputs of the prediction (CaMKII-PP2B balance) were subdivided into two classes based on whether the following constraint were fulfilled or not:

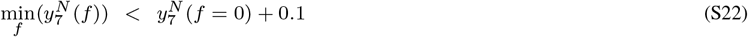

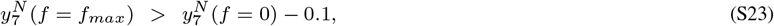

where *f* is the Ca-frequency and *f*_*max*_ is the highest frequency used in the simulation (i.e. 25 Hz).

### S4 Characteristics of the posterior distribution

A summary of the characteristics of the marginal posterior distributions are given in Table S4 for the free parameters and for the thermo-constrained parameters in Table S5. Histograms of the marginal distributions are given in Figure S1.

**Table S4.** Arithmetic mean, credibility interval and default parameter value on a log10-scale for the free *K*_*d*_-parameters and kautMax based on the marginal posterior distributions.

| Parameter | Mean | Credibility interval (95%) | Default value | Bimodal |
| --- | --- | --- | --- | --- |
| $K_d^*CaM\_Ca3^*Ca$ | 3.0 | (1.5 , 4.5) | 4.2 | yes |
| $K_d^*CaM\_Ca2^*Ca$ | 5.6 | (4.0 , 7.1) | 4.4 | yes |
| $K_d^*CaM\_Ca1^*Ca$ | 2.0 | (0.7 , 3.2) | 3.1 | no |
| $K_d^*CaM^*Ca$ | 4.8 | (3.6 , 6.1) | 3.7 | no |
| $K_d^*CaM\_Ca4^*PP2B$ | -1.5 | (-1.9 , -1.2) | -1.6 | no |
| $K_d^*PP2B\_CaM\_Ca3^*Ca$ | 0.1 | (-0.8 , 1.5) | 1.8 | no |
| $K_d^*PP2B\_CaM\_Ca2^*Ca$ | 3.8 | (1.1 , 5.3) | 2.9 | no |
| $K_d^*PP2B\_CaM\_Ca1^*Ca$ | 2.9 | (0.8 , 4.3) | 1.8 | no |
| $K_d^*PP2B\_CaM^*Ca$ | 2.7 | (0.7 , 4.9) | 2.8 | no |
| $K_d^*CaM\_Ca4^*CaMKII$ | 0.4 | (-1.2 , 1.4) | 1.7 | yes |
| $K_d^*CaMKII\_CaM\_Ca3^*Ca$ | 1.7 | (0.4 , 3.4) | 3.3 | no |
| $K_d^*CaMKII\_CaM\_Ca2^*Ca$ | 2.0 | (0.6 , 4.4) | 3.6 | no |
| $K_d^*CaMKII\_CaM\_Ca1^*Ca$ | 3.7 | (1.0 , 6.6) | 3.7 | no |
| $K_d^*CaMKII\_CaM^*Ca$ | 4.2 | (1.3 , 6.2) | 3.4 | no |
| $K_d^*pCaMKII\_CaM\_Ca3^*Ca$ | 1.6 | (0.4 , 3.2) | 3.3 | no |
| $K_d^*pCaMKII\_CaM\_Ca2^*Ca$ | 3.5 | (1.3 , 5.6) | 3.6 | no |
| $K_d^*pCaMKII\_CaM\_Ca1^*Ca$ | 4.4 | (2.0 , 6.3) | 3.7 | no |
| $K_d^*pCaMKII\_CaM^*Ca$ | 4.3 | (1.3 , 6.1) | 3.4 | no |
| $K_d^*CaM\_Ca4^*pCaMKII$ | 0.1 | (-2.7 , 1.8) | -0.1 | no |
| kautMax | 0.3 | (-1.3 , 4.7) | 1.7 | yes |

**Table S5.** Arithmetic mean, credibility interval and default parameter value for the thermo-constrained *K*_*d*_-parameters based on the marginal posterior distributions and rules applied within our model.

| Parameter | Mean | Credibility interval (95%) | Default value |
| --- | --- | --- | --- |
| $K_d^*CaM\_Ca3^*PP2B$ | 1.4 | (-1.0 , 3.5) | 0.8 |
| $K_d^*CaM\_Ca2^*PP2B$ | 3.2 | (2.2 , 4.9) | 2.3 |
| $K_d^*CaM\_Ca1^*PP2B$ | 2.3 | (-0.3 , 5.0) | 3.5 |
| $K_d^*CaM^*PP2B$ | 4.4 | (4.1 , 4.7) | 4.4 |
| $K_d^*CaM\_Ca3^*CaMKII$ | 1.7 | (-1.5 , 4.7) | 2.6 |
| $K_d^*CaM\_Ca2^*CaMKII$ | 5.3 | (3.3 , 7.2) | 3.4 |
| $K_d^*CaM\_Ca1^*CaMKII$ | 3.5 | (-1.2 , 7.7) | 2.7 |
| $K_d^*CaM^*CaMKII$ | 4.1 | (-0.7 , 8.8) | 3.0 |
| $K_d^*CaM\_Ca3^*pCaMKII$ | 1.5 | (-2.7 , 5.1) | 0.8 |
| $K_d^*CaM\_Ca2^*pCaMKII$ | 3.5 | (-0.5 , 7.5) | 1.5 |
| $K_d^*CaM\_Ca1^*pCaMKII$ | 1.2 | (-3.0 , 5.3) | 0.9 |
| $K_d^*CaM^*pCaMKII$ | 1.6 | (-1.8 , 5.8) | 1.2 |

### S5 Analytical equilibrium model reduction

Analytical solutions were obtained for phenotypes 1-4 (cf. Table S2) as follows: We note that these subsystems only consist of reversible reactions of the form *A* + *B* ⇌ C, and hence all the reaction fluxes for these subsystems will be of the form *k*_*f*_[*A*][*B*] – *k*_*r*_[*C*]. At equilibrium, all the reaction fluxes are zero, and so, we can solve the equations by writing them as *k*_*f*_[*A*][*B*] = *k*_*r*_[*C*], and then taking logarithms on both sides, giving log [*A*] + log[*B*] — log[*C*] = log(*K*_d_). Note that these equations are linear in the logarithm of the species concentrations. The number of such equations is the same as the number of reactions in the subsystem, and note that different species concentrations will come in as [*A*], [*B*] and [*C*] in the equation above. To keep track of which species are the reactants and the product in this system, we make use of the stoichiometry matrix. We number the species and reactions of the subsystem that we are considering, according to some order, and ignore other reactions and species that do not occur in this subsystem. We denote the unknown equilibrium concentrations by the column vector *X,* and the equilibrium constants by the column vector *K*_*d*_. The entries of the *K*_*d*_ vector will in this section be denoted by **K**d1, **K**d2,…, where the number of the parameter coincides with the ID of the particular reaction (as given in Table S2). The stoichiometry matrix *N* is defined by its entries

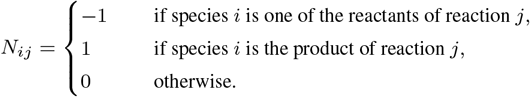

The system of equations that we need to solve can then be written in matrix form as

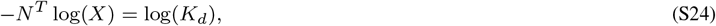

where log(*X*), log *K*_*d*_ denotes the column vector consisting of the logarithms of the entries of *X*, *K*_*d*_, respectively. This system has nontrivial solutions if and only if the vector 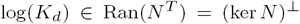, where Ran(*N*^*T*^) and ker(*N*) denote the range of *N*^*T*^ and kernel (null space) of *N,* respectively. Here, ⊥ denotes the orthogonal complement. The general solution of this system is found as the sum of the general solution to the corresponding homogeneous equation and a particular solution of the inhomogeneous system. Thus, the number of free parameters in the general solution is the same as the dimension of the null space of *N*^*T*^. Uniqueness of solutions will only be obtained when the conservation laws are taken into account. These conservation laws can also be determined from the kernel of *N*^*T*^. The conservation laws are of the form *C^T^X* = totalamount, where *C*^*T*^ is a fixed row vector of nonnegative integers, of the same length as the number of species in the subsystem. The vector *C* can be found as a basis vector of the null space of *N*^*T*^.Now we will find the explicit solutions for the subsystems.

#### S5.1 Explicit solutions for the subsystem of phenotype 1

The species that are present in this subsystem are CaM, CaM_Ca1, CaM_Ca2, CaM_Ca3, CaM_Ca4. We denote the equilibrium concentrations of these species with *X*_1_,…, *X*_5_ (with the same order as above). The stoichiometry matrix *N* is given by

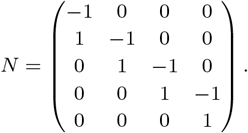

The system (S24) has nontrivial solutions for all values of the *K*_*d*_ parameters, since ker(*N*) = {0}. A basis for the null space of *N*^*T*^ is *C*_1_:= {(1, 1, 1, 1, 1)^*T*^}. As a particular solution of the system (S24), we take

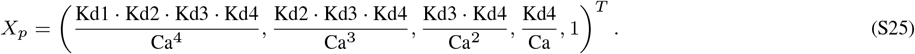

Since the nullspace of *N*^*T*^ is one-dimensional, and spanned by *C*_1_, we obtain the general solution of (S24) as

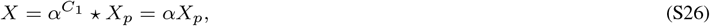

where * denotes the Hadamard (i.e. pointwise) product and 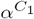 denotes the vector which is formed by raising the number *α* to each separate entry of *C*_1_. To determine α, we invoke the conservation law 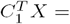 totalCaM, and obtain

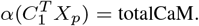

Solving this equation for *α*, and substituting the obtained expression for *α* into (S26), we obtain the equilibrium solution

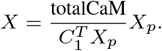

Finally, we compute the output for phenotype 1 as

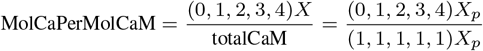

with *X*_*p*_ as in (S25).

#### S5.2 Phenotype 2

Only reaction 5 is active in this subsystem, and the equilibrium concentrations are [**CaM**] =: *X*_1_, [**PP2B**] =: *X*_2_, and [**PP2B_CaM**] =: *X*_3_. The stoichiometry matrix is *N* = ( –1, –1, 1)^*T*^. The system (S24) has nontrivial solutions for all values of *K*_*d*_ since ker(*N*) = {0}. We find a particular solution (S24) as

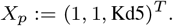

The kernel of *N*^*T*^ is spanned by *C*_1_:= (1, 0, 1)^*T*^ and *C*_2_:= (0, 1, 1)^*T*^. There are two conservation laws: 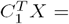 totalCaM and 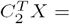 totalPP2B. The general equilibrium solution is therefore of the form

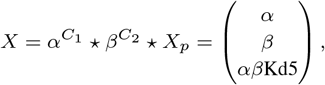

and from the conserved quantities we obtain the system of equations

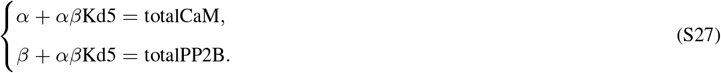

This system is reduced to a quadratic equation in *α* (by solving the second equation for *β* and substituting the solution into the first equation). The quadratic equation has one positive and one negative root, and since *α* = [CaM] cannot be negative, only the positive root is relevant. This way we obtain

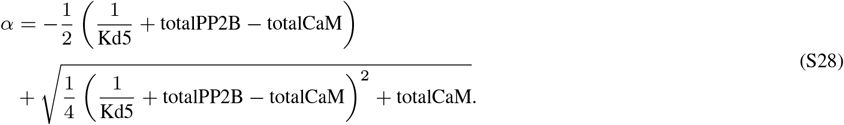

Finally, the output MolCaMPerMolPP2B is computed as

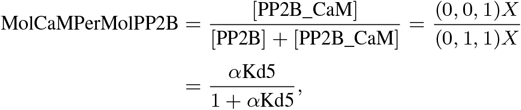

with *α* as in (S28).

#### S5.3 Phenotypes 3 and 4

The subsystems of phenotype 3 and 4 are the same. The only difference is the total amount of CaM. The subsystem has 11 species, 13 reactions and 2 conservation laws. The computations are very similar to those of phenotype 2, although the formulas are longer. The species concentrations are denoted by *X*_1_:= [CaM], *X*_2_:= [CaM_Cal], *X*_3_:= [CaM_Ca2], *X*_4_:= [CaM_Ca3], *X*_5_:= [CaM_Ca4], *X*_6_:= [PP2B], *X*_7_:= [PP2B_CaM], *X*_8_:= [PP2B_CaM_Cal], *X*_9_:= [PP2B_CaM_Ca2], *X*_10_:= [PP2B_CaM_Ca3], *X*_11_:= [PP2B_CaM_Ca4]. The nullspace of *N* has dimension 4, and it is spanned by

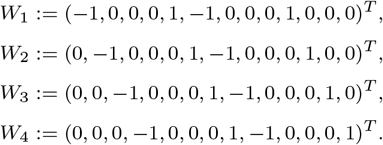

The system (S24) has a nontrivial solution if and only if log(*K*_*d*_) ∊ ker(*N*)^⊥^, i.e. if and only if log(*K*_*d*_) is orthogonal to *W*_*j, j*_ = 1,…, 4. These conditions read

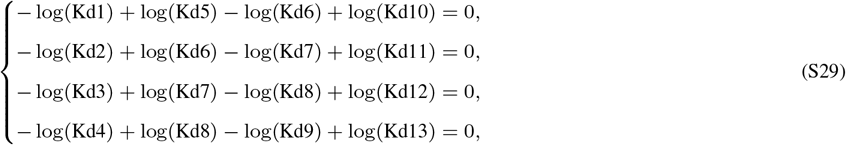

which is equivalent to the Wegscheider conditions for this subsystem (cf. Table S3).

The null space of *N*^*T*^ has dimension 2, and it is spanned by

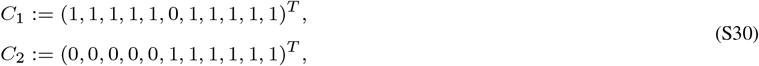

giving rise to the conservation laws *C*_1_*X* = totalCaM and *C*_2_*X* = totalPP2B. A particular solution of (S24) is given by

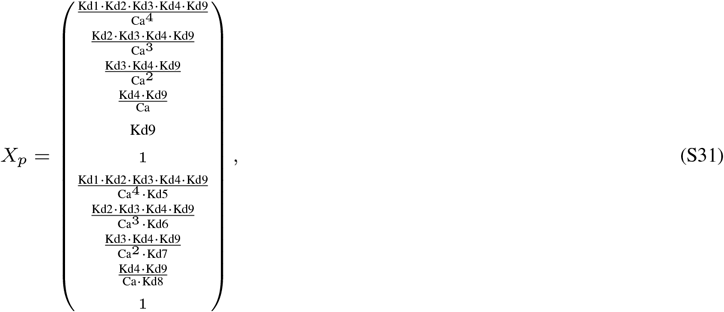

and the general solution is given by

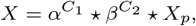

Where *C*_1_ and *C*_2_ are given by (S30) and *X*_*p*_ is as in (S31). The conservation laws give rise to the system

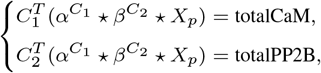

or equivalently,

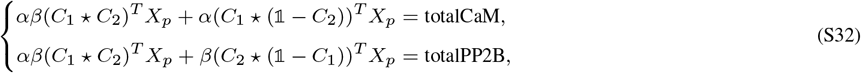

where 𝟙 is a column vector of length 11 where all entries are 1. The system (S32) can be simplified further by noting that 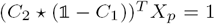, and can be solved in the same way as (S27), leading to the quadratic equation

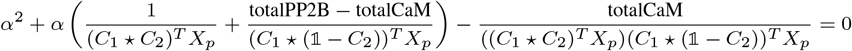

in *α* with one positive and one negative root. Again, it is only the positive root that is valid, and so

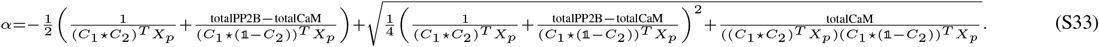

The output for both phenotypes 3 and 4 is

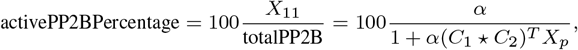

with *α* given by (S33), and (*C*_1_ * *C*_2_)^*T*^ *X*_*p*_ is the sum of the last five entries of *X*_*p*_, with *X*_*p*_ given by (S31).

#### S5.4 Phenotypes 5 and 6

Since the subsystems contains a reaction which is neither elementary nor irreversible, the method that was used in Sections S5.1–S5.3 is not applicable. For this reason, we have simulated the ODE systems instead of using analytical equilibrium solutions when computing these outputs.

**Fig. S1.**
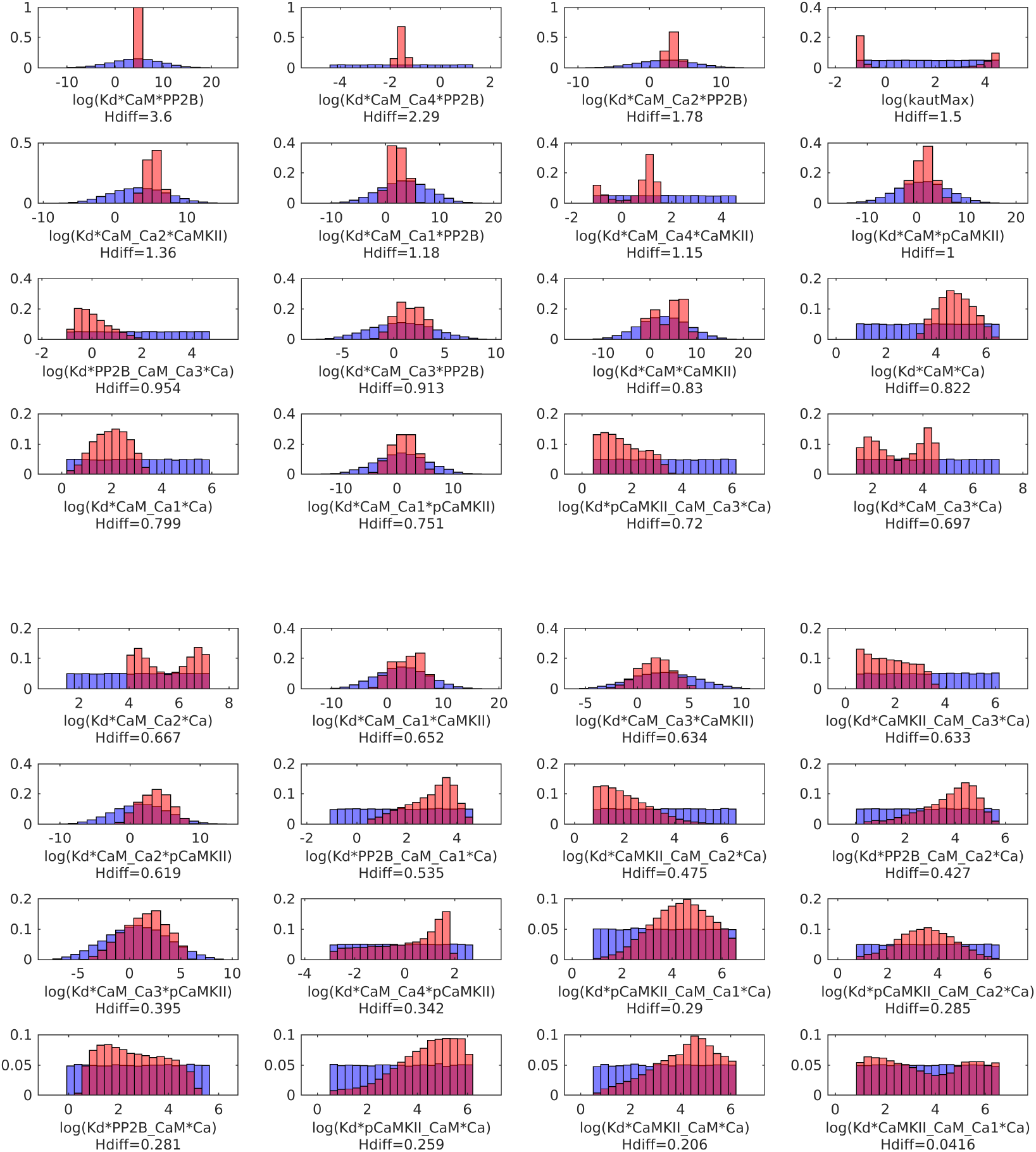
Marginal prior and posterior distributions for the model parameters sorted by reduction in entropy (H_diff_). Normalized sample histograms from the prior (blue) and posterior (red) distributions of the free and thermodynamically constrained K_D_ parameters of the model. The prior of the free parameters correspond to a sample from a log-uniform distribution centered around the default parameter values, while the priors of the thermodynamically constrained parameters are the same samples transformed by the constraint rules given in Table S3.

**Fig. S2.**
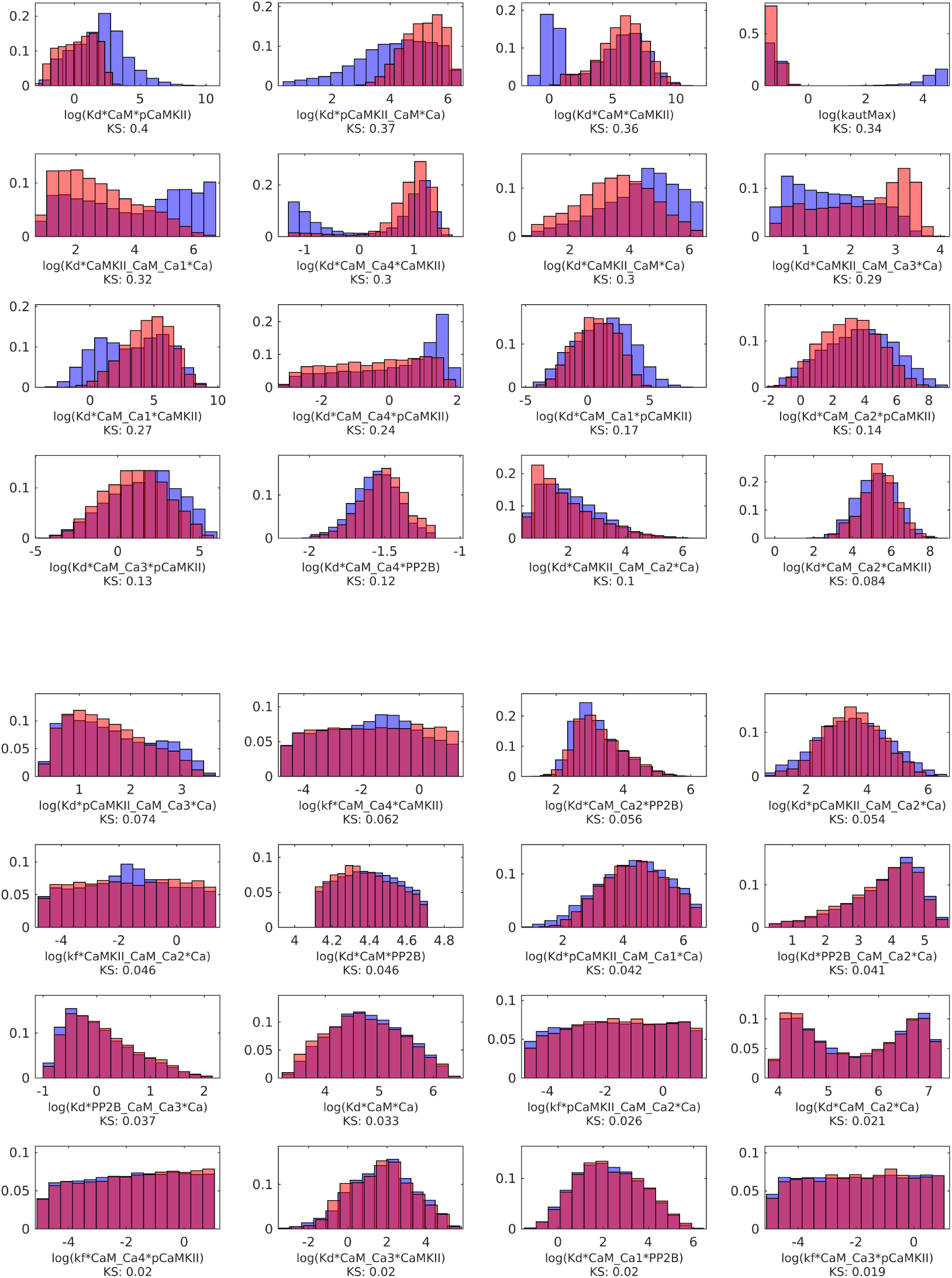
Normalized histograms describing the marginal posterior distributions of the model parameters subdivided into two classes depending on the behaviour of the output function corresponding to phenotype 5. The colors corresponds to the classes defined in figure 7 blue=monotonic (corresponding to figure 7 top, left), red=non-monotonic (corresponding to figure 7 top, right). The parameter histograms are sorted according to the Kolmogorov-Smirnov test statistica.

**Fig. S3.**
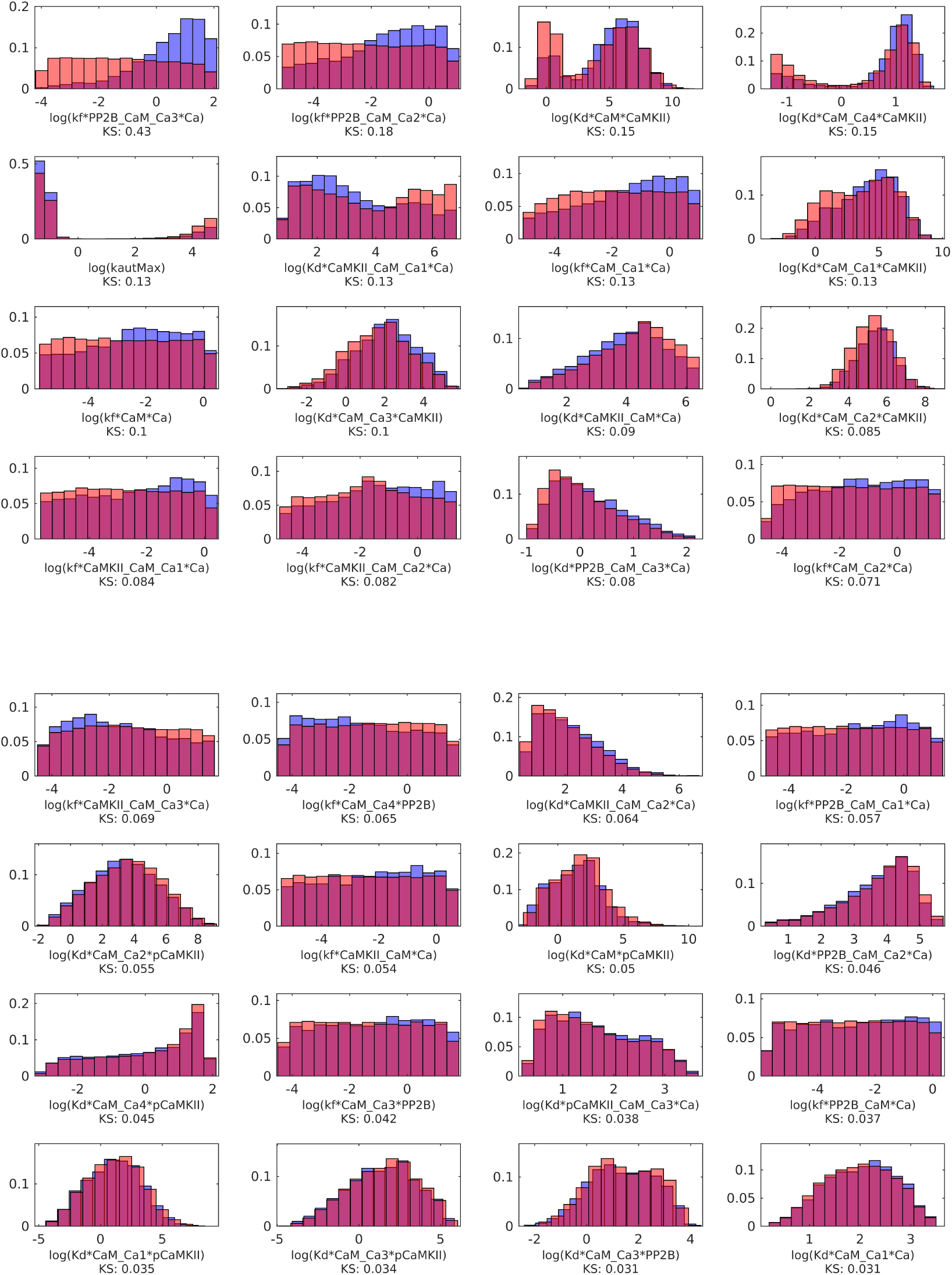
Normalized histograms describing the marginal posterior distributions of the model parameters subdivided into two classes depending on the behaviour of the output function corresponding to the prediction. The colors corresponds to the classes defined in figure 8, blue=LTD-LTP (corresponding to figure 8 top, left), red=non-LTD-LTP (corresponding to figure 8 top, right). The parameter histograms are sorted according to the Kolmogorov-Smirnov test statistica.

**Fig. S4.**
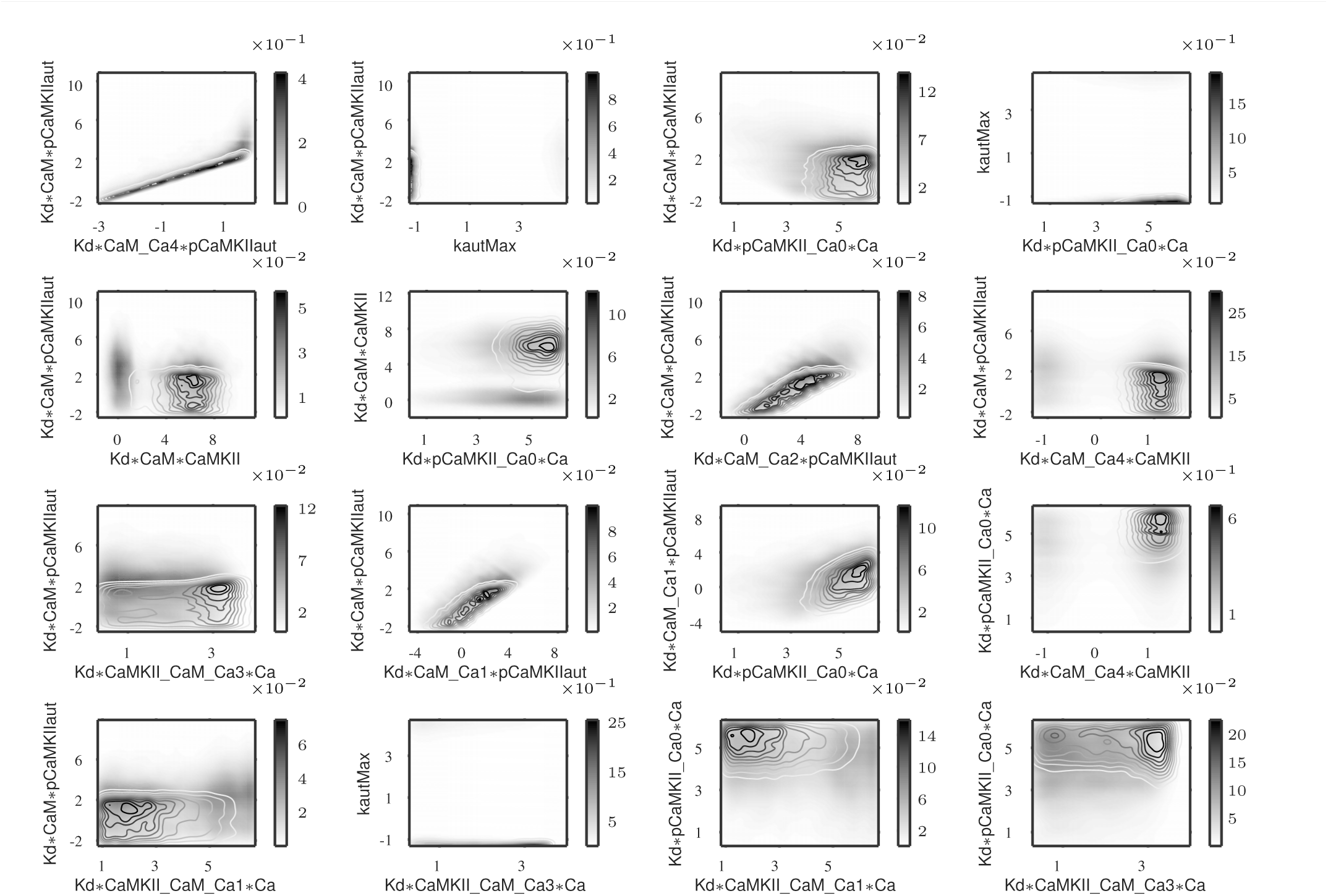
The outputs corresponding to phenotype 5 were divided into two classes, monotonic behaviour of the observed output function and non-monotonic behaviour. The subsamples corresponding to these two classes were further investigated via pairwise projections to find the pairs that cause these different behaviours. The density of parameters resulting in monotonic output behavior are represented by color shading and the parameters resulting in non-monotonic behaviour are shown as contour plots using the same color scheme. That way the lines are only visible if there is a difference in the two projected densities. If the densities are very different then the parameters pair is important for this classification. We calculated the Kullback Leibler Divergence (KLD) for each pair; this picture shows the top 16 pairs with highest KLD values.

**Fig. S5.**
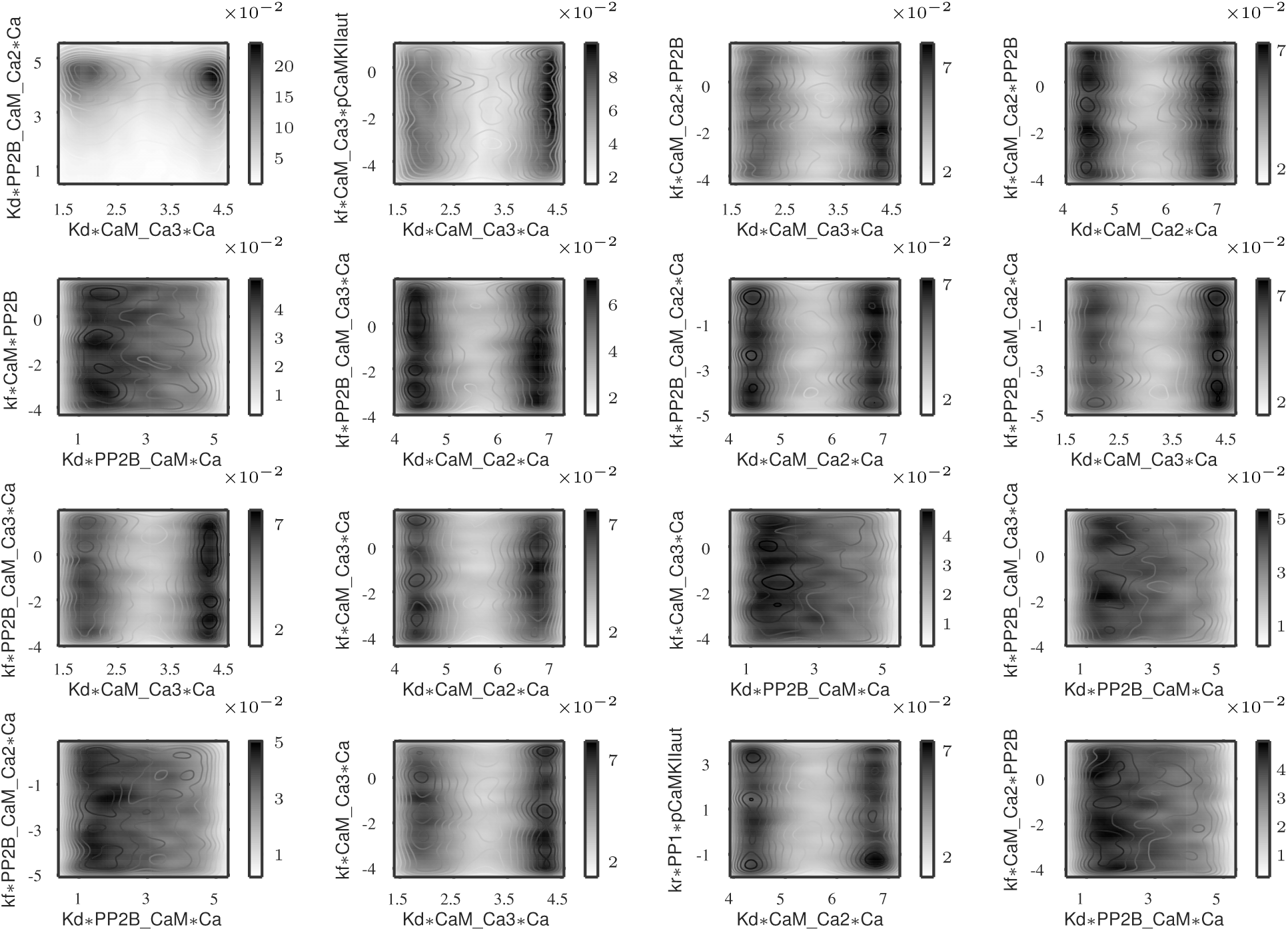
For illustration purposes, this figure shows the corresponding bottom 16 parameter pairs, ranked by KLD score. The figure was produced in the same fashion as Figure S4. The contour lines depicting parameters with resulting non-monotonic output behaviour and the color shading (for monotonic behaviour) cannot be distinguished because the densities are so similar.

**Fig. S6.**
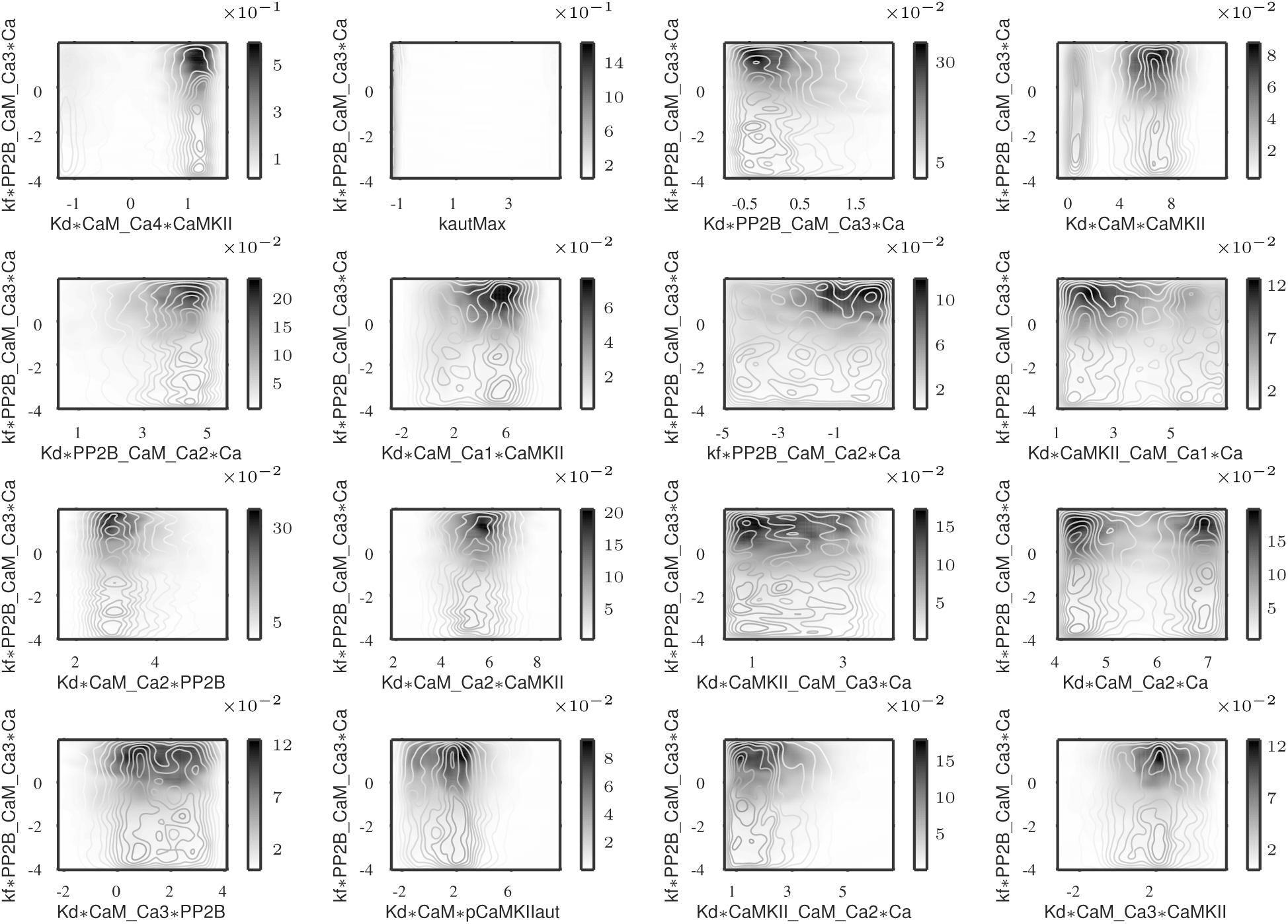
The outputs corresponding to the prediction were divided into two classes: both LTP and LTD behaviour is observed (depending on Ca frequency) or only one of those behaviours is observed regardless of Ca frequency variation. The subsamples corresponding to these two classes were further investigated via pairwise projections to find the pairs that cause these different behaviours: color shading shows the density responsible for output behaviour with both LTP and LTD, while the contour lines indicate the subsamples denity without this feature. If the two projected probability densities are very different then the parameter pair is important for this classification, in such cases the lines can be seen clearly, they are colored using the same colormap. We calculated the Kullback Leibler Divergence (KLD) for each pair; this picture shows the top 16 pairs with highest KLD values.

**Fig. S7.**
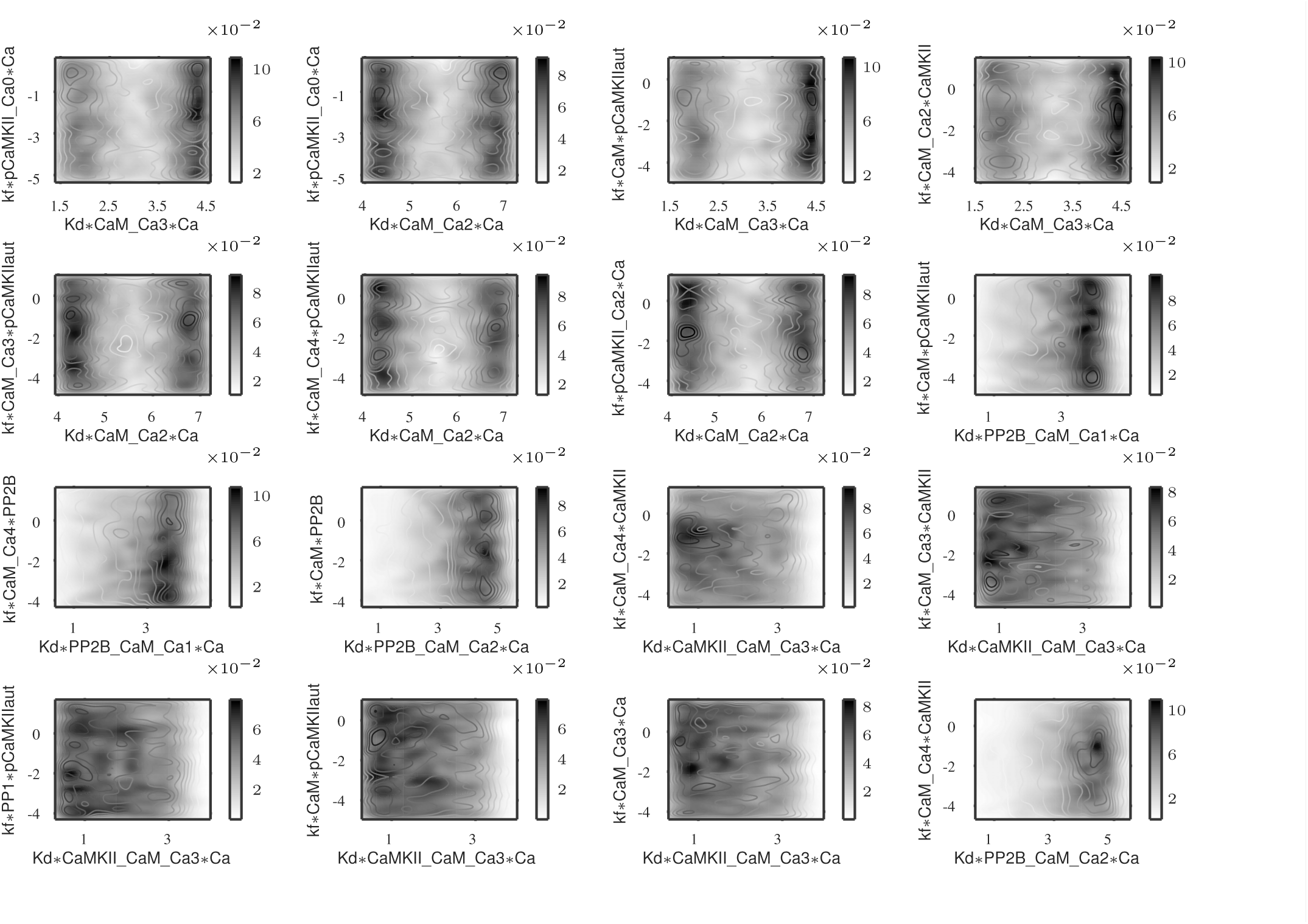
For illustration purposes, this figure shows the corresponding bottom 16 parameter pairs, with lowest KLD values. It was produced in the same fashion as Figure S6 and shows no discernible difference between the two classes for each parameter pair. The color shading indicates the density of parameter vectors that generate both LTP and LTD behaviours upon Ca frequency variation while the contour lines represent the complement of that sample (parameters not leading to both LTP and LTD in the output).

